# Modulators of epithelial-mesenchymal transitions in prostate cancer: potential for novel therapeutics development

**DOI:** 10.1101/2025.07.23.666287

**Authors:** Jinxia Zheng, Ling Li, Holly Russell, Duygu Duzgun, Sebastian Oltean

## Abstract

Research in the area of hallmarks of cancer has opened the possibility of designing novel therapeutic interventions based on modulating cancer properties. Interestingly, the epithelial-mesenchymal transition (EMT), important in both tumour growth and metastasis, has not been targeted in prostate cancer (PCa). Previously, in a repositioning screen, three new chemicals that modulate EMT in PCa (named LLSOs) have been found. This study aims to investigate how they work mechanistically as well as the effect of these LLSOs on the properties of PCa cells both *in vitro* and *in vivo*.

Although LLSOs exhibited varying effects on different cancer cell lines, all of them modulated EMT by inhibiting the migration of PCa cells. LLSO3 also exhibited a significant inhibitory effect on tumour growth *in vivo*, concomitant with an increase in the expression levels of E-cadherin. Further investigation was conducted to obtain mechanistic insights into the LLSO compounds mode of action.

In conclusion, our work strongly supports that LLSO compounds may become important candidates for targeted therapeutics in prostate cancer.

## 1. Introduction

Prostate cancer (PCa) is the most commonly diagnosed cancer in men in the Western world and is the second most common cancer in men globally [1–3]. In UK, more than 50,000 men are diagnosed with this disease, and more than 12,000 men die from PCa every year. One in eight men will be diagnosed with prostate cancer in their lifetime, and around 510,000 men are living with or after PCa in UK[4, 5]. While recent progress has been made in the last years in the treatment and management of PCa, considering the high incidence and high number of deaths, much effort is still required to find new targeted treatments.

More than 20 years ago, Hanahan and Weinberg developed an excellent framework to organize and understand the myriad of cancer properties – the hallmarks of cancer[6]. This has been very useful in cancer biology in the search for key targets and therapies. Despite its central role in cancer progression, epithelial– mesenchymal transition (EMT) remains a hallmark of cancer for which no effective targeted therapies exist.

EMT is the process through which epithelial cells transition to a mesenchymal phenotype, with the reverse – mesenchymal-to-epithelial transitions – also occurring. EMT is prominently active during embryogenesis, enabling cells to transition from epithelial to mesenchymal states for migration, before re-establishing epithelial characteristics to form new structures. EMT is shut down mainly in adult life with the exception of processes such as wound-healing. In addition, the EMT program regulates some immune cells’ properties. In the context of cancer, EMT and the reverse MET are thought to be some of the most important mechanisms for cancer progression and metastasis[7]. Consequently, intense efforts have been made to develop therapies to target EMT. One such example is the targeting of TGF-beta, one of the main regulators of EMT. While the approach has been quite successful in preclinical models, including inhibition of metastasis in tumour xenografts, the clinical development was abandoned at the initial stages due to the pleiotropic roles of TGF-beta in both human physiology and cancer development[8].

In previous work, we have designed a bi-fluorescent reporter based on splicing of the fibroblast growth factor receptor 2 and demonstrated that it could be used as a sensor of EMT in cancer cells[9] . More recently, using this reporter, we performed a repositioning screen using the LOPAC library to find compounds that can act as modulators of EMT. We identified three compounds (LLSO1 - NNC-55-0396 dihydrochloride; LLSO2 - Nemadipine A; LLSO3 - Naltrexone hydrochloride) and conducted preliminary validations. Here, we show that these compounds modulate EMT markers in prostate cancer cells as well as some breast cancer cell lines. Furthermore, we describe their mechanism of action and how they influence properties of cancer cells such as migration, growth or apoptosis; we also demonstrate inhibition of tumour growth *in vivo* in PCa xenografts. Therefore, our work strongly supports the use of these compounds as an effective anti-cancer therapy.

## 2. Materials and Methods

### Cell lines and cell culture

PC3 and DU145 prostate cancer cell lines were obtained from Microvascular Research Laboratories (MVRL), University of Bristol. LNCaP prostate cancer cells were purchased from the American Type Culture Collection (ATCC, Manassas, VA, USA). MDA-MB-231 and MCF-7 breast cancer cell lines; CALU-3 non-small-cell lung cancer cells; HCT-116 colorectal carcinoma cells; and SKOV-3 and A2780 ovarian cancer cells were all obtained from the Clinical and Biomedical Sciences Laboratories (CBS), University of Exeter.

PC3, DU145, LNCaP, SKOV3, and A2780 cells were maintained in RPMI 1640 (Sigma) supplemented with 10% Fetal Bovine Serum (FBS, Gibco) and 1% Penicillin streptomycin (Sigma). MCF-7, MDA-MB-231, HCT116, and CALU-3 cells were cultured in Dulbecco’s Modified Eagle’s Medium (DMEM, Sigma-Aldrich) with 10% FBS (Gibco) and 1% Penicillin streptomycin (Sigma). All cell lines were incubated under standard conditions: a humidified incubator at 37°C with 5% CO_2_.

### Western Blot

Cells were lysed with a RIPA lysis and extraction buffer (Thermofisher) containing 1% Halt proteinase inhibitor cocktail (Thermofisher). Protein samples were denatured in Laemmli sample buffer (Bio-Rad) with 2-Mercaptoethanol (Bio-Rad), total proteins were loaded onto TGX stain-free protein gels (Bio-Rad), for Western transfer to PVDF membranes (Bio-Rad) and immunoblotting. Following this, membranes were blocked with 5% BSA in 50 mm Tris-HCl (pH 7.4), 150 mm NaCl, and 0.3% Tween 20 for one hour, immunoblots were probed overnight at 4°C with the following primary antibodies: anti-E-cadherin (BD Biosciences), anti-N-cadherin (Abcam), and anti-Twist (GeneTex). After washing, membranes were incubated with a fluorescent secondary antibody (Li-Cor) for an hour and imaged using a Licor imager. Blots were analysed by quantitative densitometry using Bio-Rad ImageLab software.

### Immunofluorescence

Cells were seeded onto coverslips in 6-well plates and allowed to adhere for 24 hours. Following treatment with either DMSO or various LLSOs for 48 hours, cells were fixed with ice-cold methanol for 10 minutes and permeabilized with 0.05% Triton X-100 (Thermofisher) in PBS for 10 minutes, followed by permeabilization. Nonspecific binding sites were blocked with PBS containing 1% Bovine Serum Albumin (BSA, Sigma-Aldrich) and 5% normal goat serum (Abcam) for one hour at room temperature. Cells were then incubated with either Alexa Fluro 488 Goat anti-Mouse IgG (Invitrogen) as the isotype control or anti-E-cadherin (BD Biosciences), and anti-N-cadherin (Abcam) primary antibodies in PBS with 1% BSA overnight at 4 °C. After three washes with PBS, cells were incubated with Alexa Fluor 488 goat anti-mouse secondary antibody (Invitrogen) for 90 minutes and mounted in VECTASHIELD with DAPI (VECTASHIELD® Antifade Mounting Medium with DAPI, Vector Labs). Fluorescence images were acquired using Leica DM4000 LED fluorescence microscopy and image analysis was performed using ImageJ software.

### Migration assay (Boyden chamber assay)

Cells were pre-treated with either DMSO or LLSOs for 48 hours. LNCaP cells were subsequently serum-starved overnight in serum-free medium, while DU145 cells were cultured without starvation due to their rapid migration characteristics. After treatment and cell starvation, 150,000 treated cells in serum-free medium were seeded into Millicell hanging cell culture inserts (Merck) placed in 24-well plates containing complete medium with 10% FBS as a chemoattractant. Following 24 hours of incubation, non-invasive cells that stayed in the interior of the insert were removed by gently wiping with a moistened cotton swab. Migrated cells on the outer side of the membrane were fixed and stained with DAPI. Cell counts were obtained from three random fields per insert using fluorescence microscopy. Data were analysed by one-way ANOVA with Dunnett’s multiple comparisons test using GraphPad Prism 9.

### Apoptosis assay (JC-10 mitochondrial membrane potential assay)

Cells were seeded in 90μl of media per well in a black-walled, clear-bottom 96-well culture microplate and cultured overnight at 37℃ and 5% CO2. Following treatment with DMSO or various concentrations of LLSOs for 48 hours, the JC-10 assay kit (Abcam) was used according to the manufacturer’s protocol. Fluorescence intensity was measured using the SpectraMax M2e Multi-mode Plate Reader with the following settings: Ex/Em = 490/525 nm (cut off at 515 nm) and 540/590 nm (cut off at 570 nm) for ratio analysis.

### Population growth curve of the combination treatment

50,000 of PC3 cells or 70,000 of DU145 cells were seeded per well in 24-well plates. After 24 hours, cells were treated with DMSO as a control, 5µM LLSO1, 10µM LLSO2, 10µM LLSO3, 5µM LLSO1 + 10µM LLSO2, 5µM LLSO1 + 10µM LLSO3, 10µM LLSO2 + 10µM LLSO3 or 5µM LLSO1 + 10µM LLSO2 + 10µM LLSO3. Cells were counted every 24 hours for a total of 120 hours, with media and treatments refreshed every 48 hours. Data were analysed with GraphPad Prism 9 and tested by two-way ANOVA with Dunnett’s multiple comparisons test.

### Mice xenografts

All the *in vivo* experiments were approved by the University of Exeter Animal Welfare and Ethical Review Body (AWERB) and conducted in accordance with UK legislation. CD-1 nude mice (Charles River Laboratories, Wilmington, MA, USA) were subcutaneously injected with 3 million PC3 cells in 100 µL PBS (n = 12). Once tumours reached 3 mm in diameter, mice were randomised into two groups (n = 6/group) and injected intraperitoneally three times weekly with either saline (control) or 10 µM LLSO3. Tumour size was measured three times weekly using calipers, and volume was calculated as [(length + width)/2] × length × width. Once the tumour size reached the allowed endpoint with a diameter of 12mm in any direction, mice were culled by cervical dislocation and the tumours were extracted. Tumours were photographed before being flash frozen in liquid nitrogen for further analysis. Statistical significance was assessed by using Two-way ANOVA through GraphPad Prism 9.

### Quantitative polymerase chain reaction (qPCR)

Total RNA was extracted by TRIzol reagent (Ambion) and total RNA (2ug) from each sample was reverse-transcribed with GoScript Reverse Transcription Mix and random primers (Promega) following the manufacturer’s instructions. Quantitative real-time PCRs (qPCR) was performed using PowerUp SYBR Green Master Mix (Thermofisher) on a LightCycler 96 Real-Time PCR System (Roche). The following oligonucleotides were used: E-cadherin F: 5’-TCCGAAGCTGCTAGTCTGAG-3’; R: 5’-CTCAAGGGAAGGGAGCTGAA-3’. The oligonucleotides used for normalisation were GAPDH F: 5’-GTCAGTGGTGGACCTGACCT-3’; R: 5’-TGACAAAGTGGTCGTTGAGG-3’.

### LC50 (MTT)

PC3 cells (10,000/well) were seeded in a 96-well plate and incubated at 37℃ and 5% CO2 for 24 hours. After cells were treated with either DMSO or different concentrations of LLSO for 48 hours, the MTT reagent (Abcam) was added per the manufacturer’s instructions, including wells without cells as background controls. Absorbance at OD=590nm was measured using a SpectraMax M2e Multi-mode Plate Reader, with blank-corrected values used to calculate viability. The data were analysed through GraphPad Prism 9 by using one-way ANOVA with Dunnett’s multiple comparisons test.

### Alamar blue assay

Cells (10,000/well) were seeded in a 96-well plate for 24 hours, then treated with either DMSO, 5μM LLSO1, 10 μM LLSO2 or 10μM LLSO3. After 48 hours of treatment, viability was assessed using AlamarBlue reagent (Thermofisher) per the manufacturer’s instructions, with fluorescence measured on a SpectraMax M2e plate reader (Molecular Devices). Data was analysed through GraphPad Prism 9 by using one-way ANOVA with Dunnett’s multiple comparisons test.

## 3. Results

### 3.1. Dose-dependent effect of LLSOs on the expression of EMT markers in prostate cancer cells

To investigate how three different LLSO compounds induce MET through modulating the expression level of EMT markers, several concentrations that are close to the LC_50_ (LC-lethal concentration) for each LLSO were tested, and the expression level of several EMT markers was analysed by western blot (**Figure1**) and immunofluorescence (**Figure2**) in prostate cancer cell lines. (The calculation of LC_50_ for each compound in PC3 prostate cancer cells is presented in **Supplementary Figures 1**, 2 and 3).

The expression of the epithelial marker E-cadherin (as assessed by Western blot) is increased, and the mesenchymal markers N-cadherin and Twist expression levels are decreased in PC3 cells incubated with the LLSO compounds (see quantifications in **Figure 1** and blots in **Supplementary Figure 4**, 5**, 6**).

**Figure 1.**
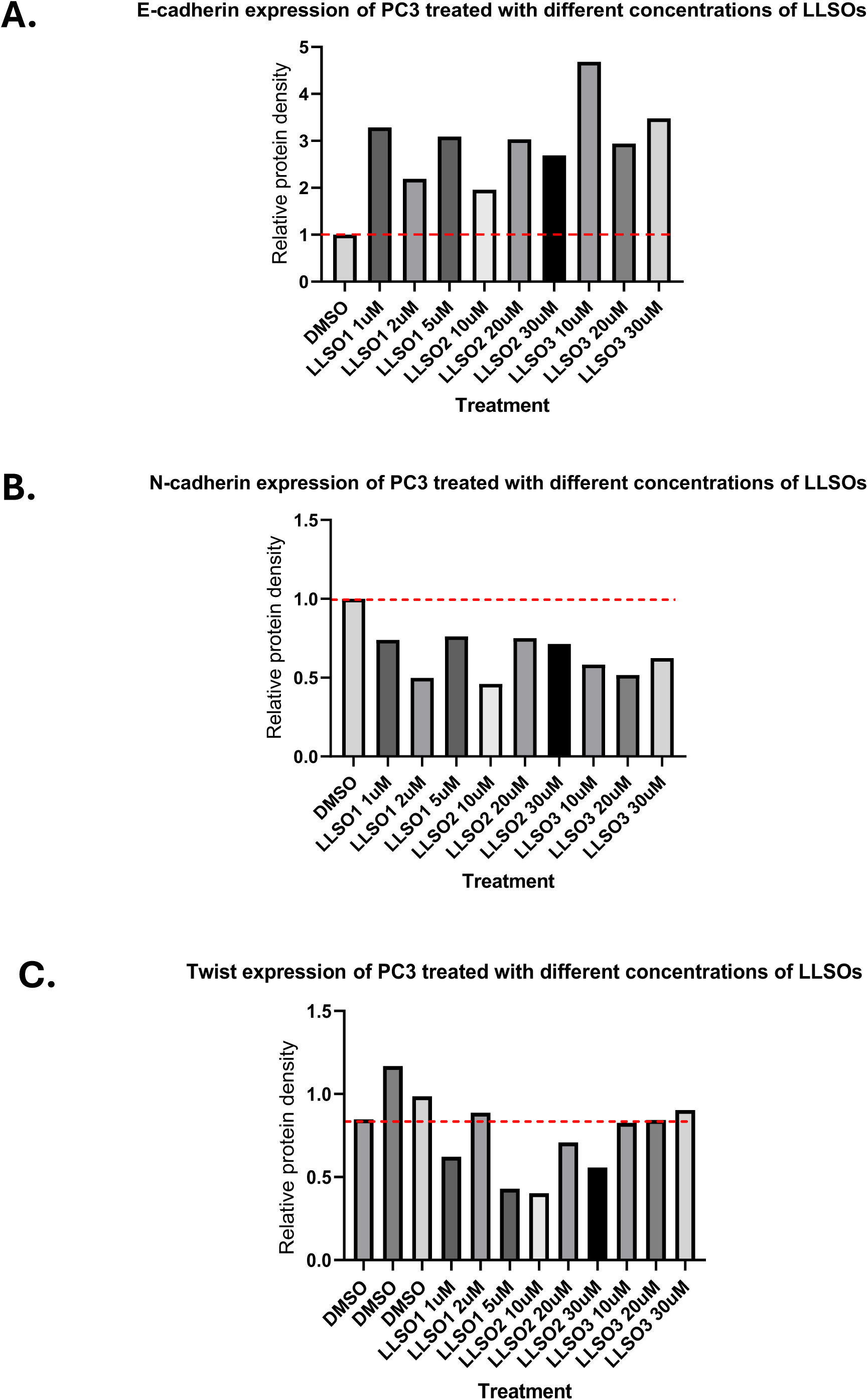
LLSOs increase the expression of E-Cadherin and decrease expression of N-cadherin and Twist in PC3 cells, as assessed by western blot. **A.** Quantification of E-Cadherin protein levels in Western blot from PC3 cells treated with DMSO (control) or different concentrations of LLSOs. All three chemicals showed increased the levels of E-cadherin expression. N=2, The experiment was repeated twice, data was analysed by one-way ANOVA. **B.** Quantification of N-Cadherin protein levels in western blot from PC3 cells treated with DMSO (control) or different concentrations of LLSOs. All three chemicals showed decreased the levels of N-cadherin expression. N=1, The experiment was done once, data was analysed by one-way ANOVA. **C.** Quantification of Twist protein levels in western blot from PC3 cells treated with DMSO (control) or different concentrations of LLSOs. All three chemicals showed decreased trend on Twist expression level. N=1, The experiment was done one time. Data was analysed by one-way ANOVA.

The expression of EMT markers was also assessed by immunofluorescence. **Figure 2** shows increase in E-cadherin fluorescence and decrease in N-cadherin when PC3 cells are treated with LLSOs. Similar results, albeit with smaller changes in expression levels, were obtained in another prostate cancer cell line – DU145 (see **Supplementary Figure 7** and 8).

**Figure 2.**
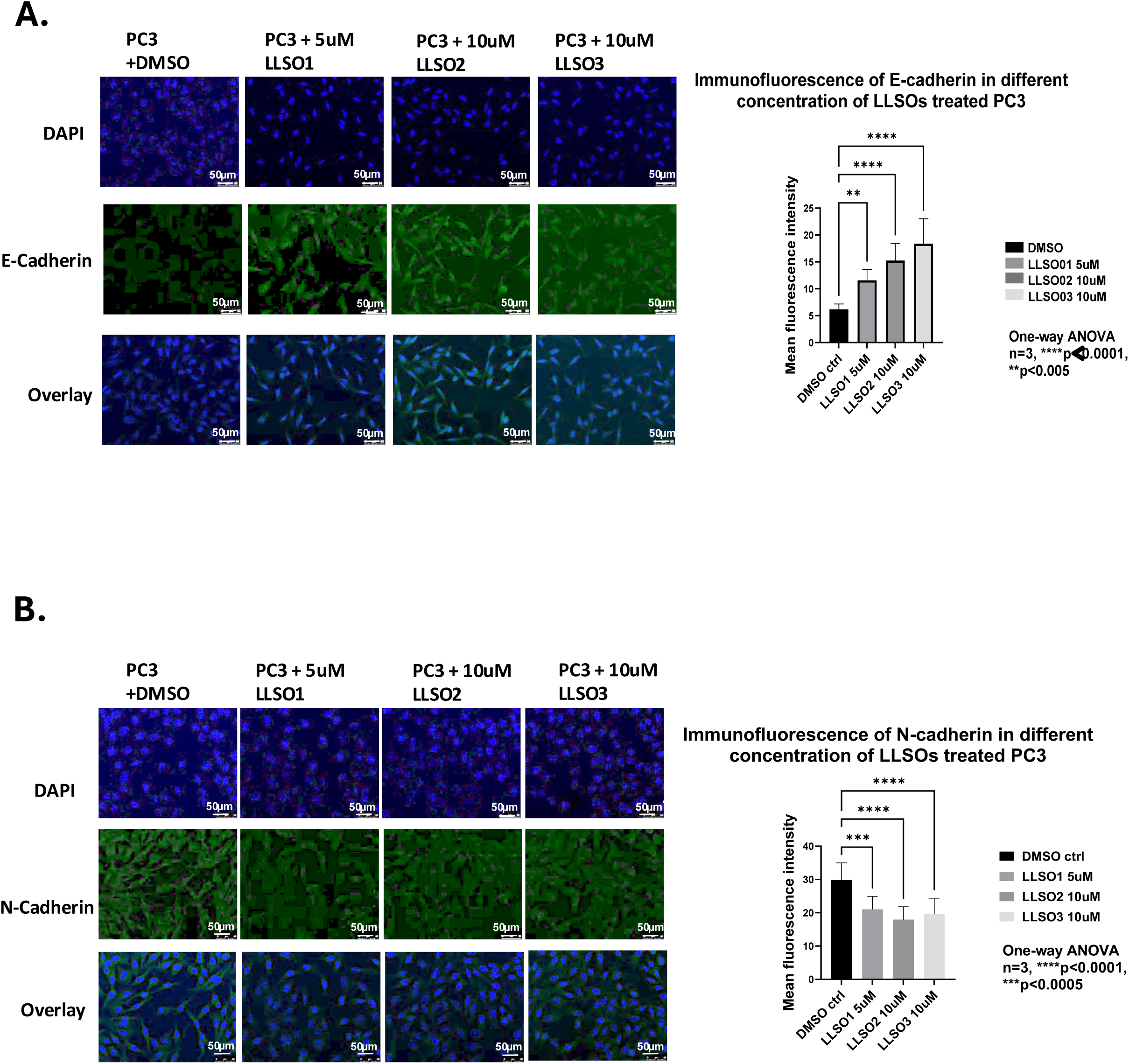
LLSOs increase the expression of E-cadherin in PC3 cells, as assessed by immunofluorescence. Immunofluorescence was performed following 48 hours of pre-treatment with DMSO or 5µM of LLSO1, 10µM of LLSO2 and 10µM of LLSO3 in PC3 cells. **A.** *Left panel* - Representative fields taken at 200x magnification; *Right panel* - Normalized E-cadherin expression of LLSO1 5µM, LLSO2 10µM, LLSO3 10µM treated PC3 cells and DMSO (as control) treated PC3 cells. n=3, Representative examples from at least 9 microscopic fields per experiment, experiments repeated 3 times with triplicate per treatment. Pictures were captured at 200x magnification, and the scale bar is 50µm. Data were analysed by one-way ANOVA **B.** *Left panel* - Representative fields taken at 200x magnification; *Right panel* - Normalized N-cadherin expression of LLSO1 5µM, LLSO2 10µM, LLSO3 10µM treated PC3 cells and DMSO (as control) treated PC3 cells. n=3, Representative examples from at least 9 microscopic fields per experiment, experiments repeated 3 times with triplicate per treatment. Pictures were captured at 200x magnification, and the scale bar is 50µm. data were analysed by one-way ANOVA

We therefore concluded that indeed the LLSO compounds are modulators of EMT at non-toxic concentrations.

### 3.2. The effects of LLSO compounds on prostate cancer cell function

We next investigated how LLSO compounds affect prostate cancer cell behaviours, both EMT-related and independent.

Previous data from our group showed that both LLSO1 and LLSO3 reduced the cell growth rate, all three LLSOs inhibited cell migration and LLSO1 decreased the proliferation rate of PC3 cells. The effect of LLSOs on LNCaP and DU145 properties *in vitro* was verified.

All three LLSOs have been shown to inhibit cell migration in PC3 cells by the Boyden chamber assay[10]. As shown in **Figure 3 A-D**, all LLSOs significantly decreased the migration rate for both LNCaP and DU145, as expected given their EMT-inhibitory activity.

**Figure 3.**
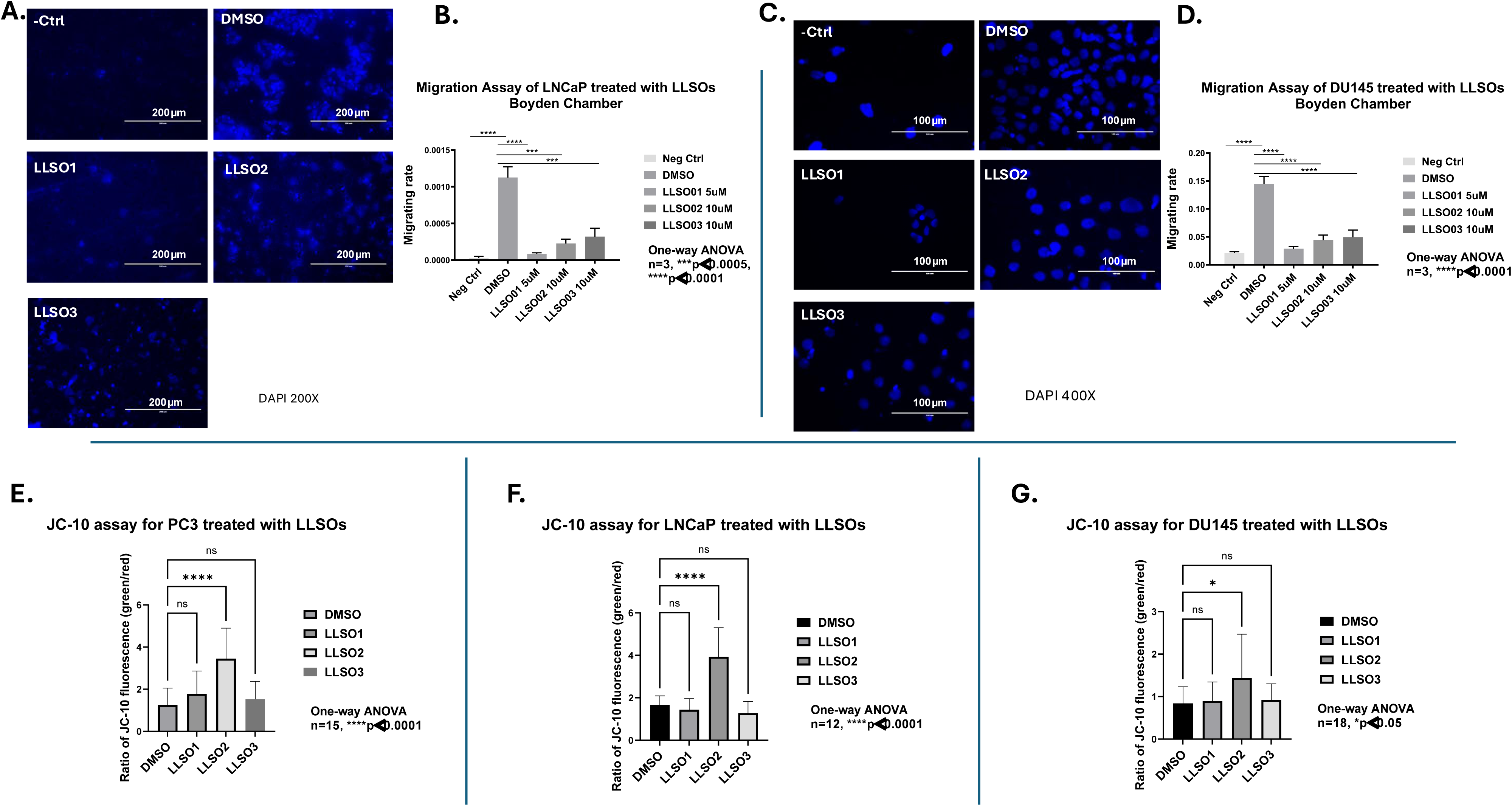
LLSOs decrease the migration rate in both LNCaP and DU145 cells and LLSO2 induces apoptosis in PC3, LNCaP, and DU145 cell lines. Boyden Chamber assay was performed following 48 hours of pre-treatment of LNCaP and DU145 cells with DMSO and LLSOs. LNCaP and DU145 cells were cultured for 24 hours in the reduced medium for starving cells. Representative images for LNCaP cells were taken at 200x magnification, and 400x magnification for DU145 cells. **A.** Representative photographs of Boyden chamber assay for either DMSO or one of the LLSOs for pre-treated LNCaP cells. LNCaP cells not starved were used as the negative control. **B.** Normalized migration rate on Boyden chamber assay of 5µM LLSO1, 10µM LLSO2, 10µM LLSO3 or DMSO treated LNCaP cells by counting cells on the Photomicrographs. Data were analysed by one-way ANOVA. experiment was repeated three times with three replication per treatment in each individual experiment, ***p<0.0005, ****p<0.0001. **C.** Representative photographs of Boyden chamber assay for either DMSO or one of the LLSOs for pre-treated DU145 cells. DU145 cells not starved were used as the negative control. **D.** Normalized migration rate on Boyden chamber assay of LLSO1 5µM, LLSO2 10µM, LLSO3 10µM and DMSO treated DU145 cells by counting cells on the Photomicrographs. Data were by one-way ANOVA, three individual experiments with three replication per treatment in each individual experiment, ****p<0.0001. **Lower panel**: JC-10 assay was performed following 48 hours pre-treatment with 5µM LLSO1, 10µM LLSO2, 10µM LLSO3 or DMSO of PC3, LNCaP and DU145 cells, as well as pre-treatment with 5mM H2O2 as the positive control. Normalized apoptosis ratio on JC-10 assay of either DMSO control or one of the three LLSOs treated (**E**) PC3, (**F**) LNCaP and (**G**) DU145 cells by using the fluorescence intensity ratio of green to red fluorescence. Data were analysed by one-way ANOVA, *p<0.05, ****p<0.0001. For PC3, three individual experiments with five replicates per treatment in each individual experiment, n=15; For LNCaP, two individual experiments with six replicates per treatment in each individual experiment, n=12; For DU145, three individual experiments with six replicates per treatment in each individual experiment, n=18.

The potential of LLSOs to induce cell apoptosis in either PC3, LNCaP or DU145 was evaluated by using JC-10 mitochondrial membrane potential assay. As shown in **Figure 3 E-G**, LLSO2 induced cell apoptosis in all cell lines, whereas both LLSO1 and LLSO3 did not affect apoptosis in any of these PCa cell lines.

The effect of LLSOs on cell growth for LNCaP and DU145 was assessed evaluating the cell growth. As shown in **Supplementary Figure 9**, 5µM LLSO1 was toxic in LNCaP cells (though this is similar to LC50 for LLSO1 in PC3 cells). The growth of LNCaP cells was inhibited by LLSO1 and all cells were killed after 120 hours of exposure to 5µM LLSO1. Both 10µM LLSO2 and 10µM LLSO3 showed a significant reduction of LNCaP growth. Similarly, 5µM LLSO1(**Supplementary Figure 10)** also decreases significantly in the growth of DU145 cells while there is no significant effect for both 10µM LLSO2 and 10µM LLSO3.

Alamar Blue assay was performed to verify the effect of LLSOs on cell viability of either PC3, LNCaP or DU145 cells. As seen in **Supplementary Figure 11**, the three LLSOs treatments showed no significant effects in both PC3 and DU145 cells, while they slightly reduced the cell viability of LNCaP cells.

### 3.3. LLSOs have cumulative effects

To determine whether LLSOs have cumulative/synergistic effects, the growth curves in PC3 and DU145 were repeated, treating the cells with combinations of various LLSOs. As seen in **Figure 4A**, all different combinations of LLSOs significantly reduced the growth of PC3 cells. Treatment with LLSO1 alone almost completely abolished growth of PC3 cells, making it difficult to determine whether combined LLSO1 + LLSO2/3 treatment had an additive effect. We also found combined treatment with LLSO2 and LLSO3 increased growth inhibition compared to LLSO3 alone. However, this growth inhibition was comparable to LLSO2 alone. Therefore, in PC3 cells, the cumulative effect of LLSOs on growth was unclear.

**Figure 4.**
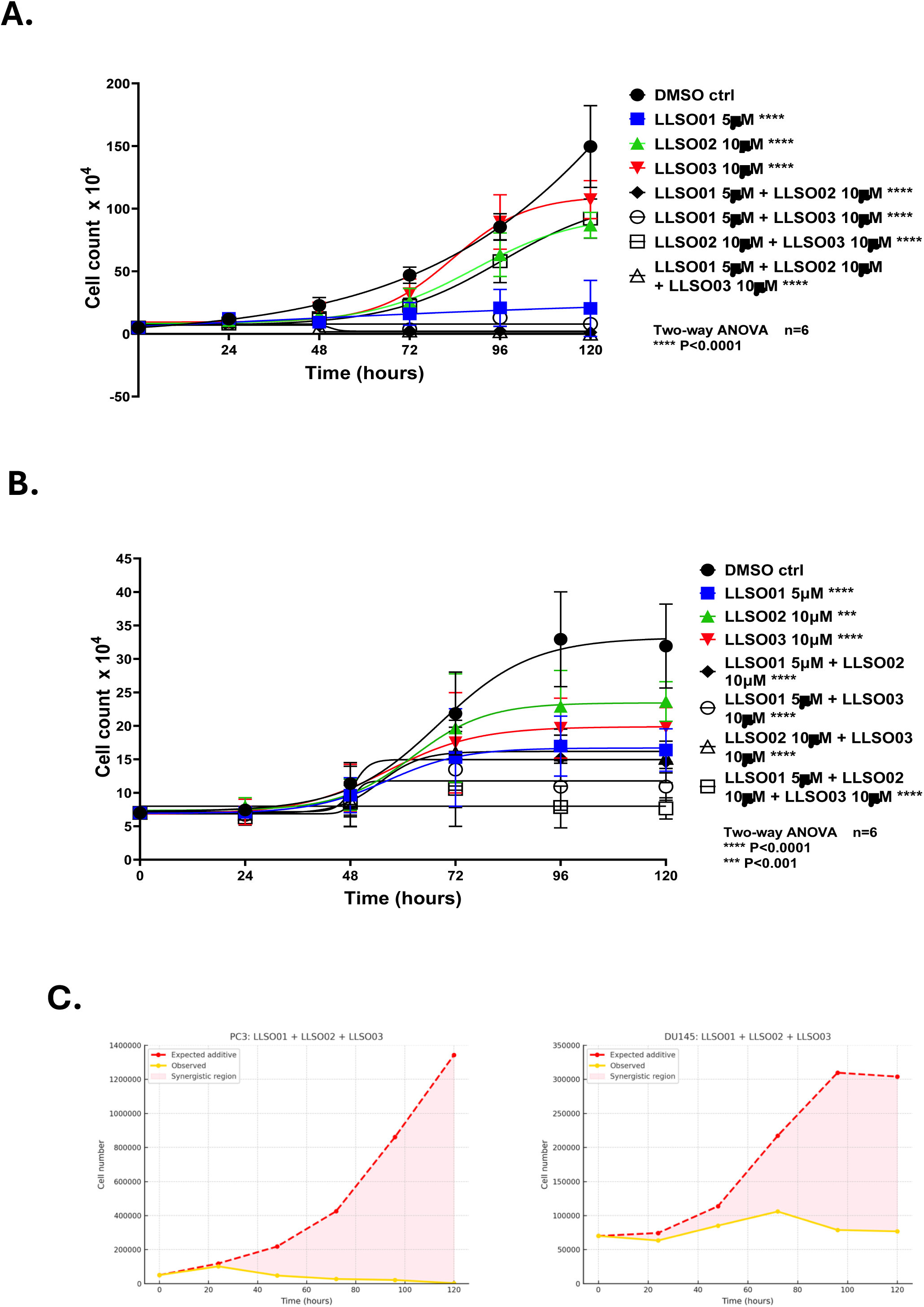
The cumulative effect of the LLSOs on the growth of PC3 and DU145. **A.** Growth curve of PC3 cells treated with a combination of LLSO or DMSO. All LLSOs treatment either on their own or in different combinations inhibited the growth of PC3 cells. Treated PC3 cells with DMSO control, 5µM LLSO1, 10µM LLSO2, 10µM LLSO3, 5µM LLSO1 + 10µM LLSO2, 5µM LLSO1 + 10µM LLSO3, 10µM LLSO2 + 10µM LLSO3 or 5µM LLSO1 + 10µM LLSO2 + 10µM LLSO3 after cells were seeded in 24-well plates for 24 hours. Medium with LLSOs treatment was refreshed every 48 hours, and cells were counted every 24 hours after 24 hours of seeding PC3 cells for a total of 120 hours. Two individual experiments with three repeats for each treatment in each individual experiment, data was analysed by two-way ANOVA. n=6, ****p<0.0001. **B.** Growth curve of DU145 cells treated with a combination of LLSO or DMSO. LLSOs had a cumulative effect on DU145 cell growth, showing greater inhibition of cell growth in any combination compared with the treatment of any of the three LLSOs alone. Treated DU145 cells with DMSO control, 5µM LLSO1, 10µM LLSO2, 10µM LLSO3, 5µM LLSO1 + 10µM LLSO2, 5µM LLSO1 + 10µM LLSO3, 10µM LLSO2 + 10µM LLSO3 or 5µM LLSO1 + 10µM LLSO2 + 10µM LLSO3 after cells were seeded in 24-well plates for 24 hours. Medium with LLSOs treatment was refreshed every 48 hours, and cells were counted every 24 hours after 24 hours of seeding DU145 cells for a total of 120 hours. Two individual experiments with three repeats for each treatment in each individual experiment, the data was analysed by two-way ANOVA. n=6, ***p<0.001, ****p<0.0001. **C.** Bliss independence analysis in both PC3 and DU145 cells showed synergy of the 3 compunds.

We also evaluated these drugs in DU145 cells (**Figure 4B**). We found treatment of DU145 cells with all three LLSO compounds simultaneously resulted in the strongest growth inhibition, completely abrogating cell proliferation. Bliss independence analysis revealed synergistic interactions among the three compounds **(Figure 4C)**, as well as between various pairs of LLSOs in both PC3 and DU145 cells **(Supplementary Figure 12).**

### 3.4. The effects of LLSO3 on tumour growth in prostate cancer xenograft models in nude mice

LLSO1 is shown to be toxic in some assays / concentrations (see above LC50 and growth curve) and LLSO2 has been previously reported[10]. We therefore evaluated the effect of LLSO3 *in vivo* in mice tumour xenografts. **Supplementary Figures 13 A and B** show the tumour growth traces in individual mice. As shown in the **Figure 5A**, 10μM LLSO3 inhibited tumour growth in PC3 xenografts in nude mice.

**Figure 5.**
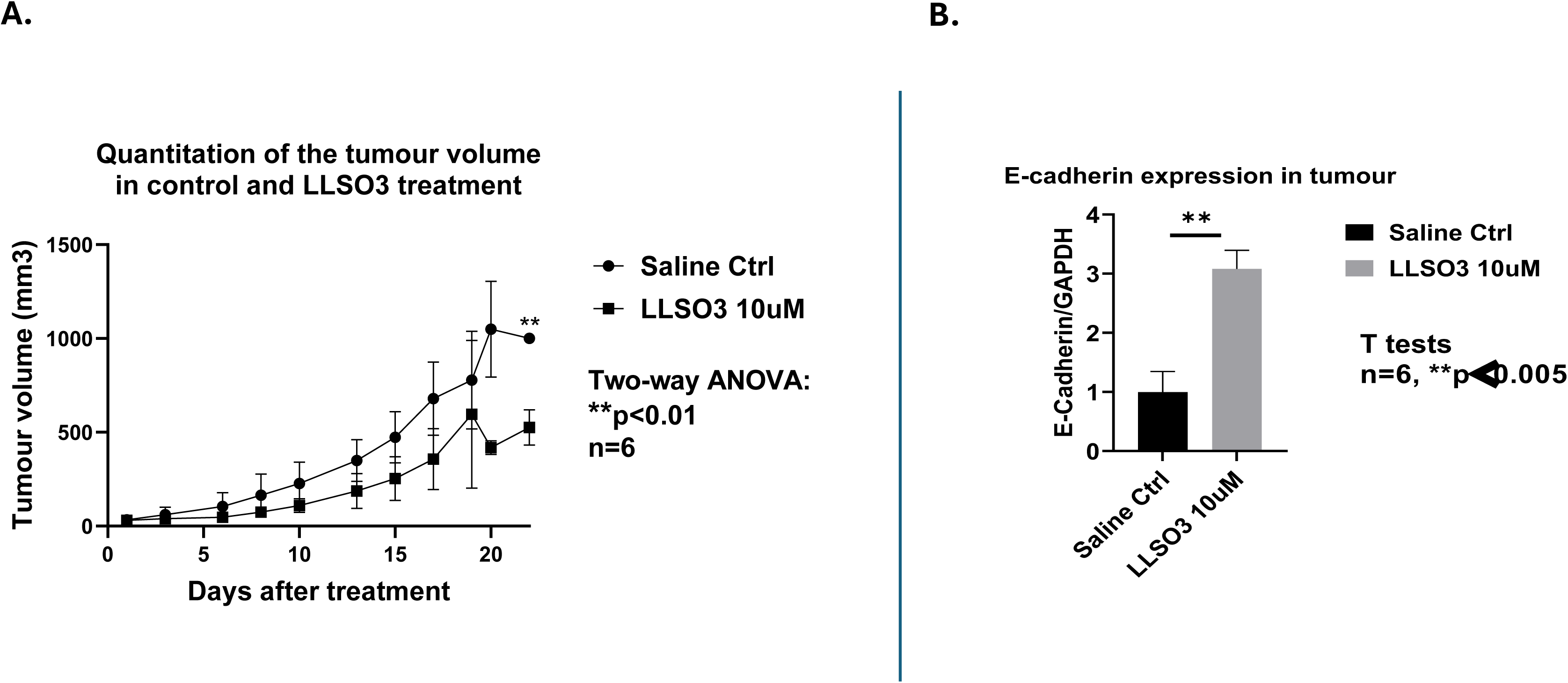
10μM LLSO3 inhibit tumour growth in PC3 xenografts in nude mice by increasing E-cadherin levels. **A.** Quantitation of the tumour volumes in saline (control) and 10μM LLSO3 treated mice at different times post-treatment. Tumour volumes were estimated using the formula ‘volume = [(length + width)/2] x length x width’. Quantitation of the tumour volumes was analysed by Two-way ANOVA using GraphPad Prism9. **B.** qRT-PCR for E-cadherin expression was performed with extracts from control and 10μM LLSO3 treated PC3 tumours. Normalization of the relative gene expression using GAPDH as the reference control. There are two repetitions within each of the 3 independent tumour samples. n=6. Results were expressed as Mean±SD, *p<0.05, **p<0.01

In order to clarify whether LLSO3 inhibit PCa tumour growth through regulation of EMT in vivo, the expression level of E-cadherin in tumour xenografts was investigated by qPCR using RNA extracted from excised tumour tissues. qPCR analysis from **Figure 5B** demonstrates that LLSO3 increases the expression level of both E-cadherin in tumours.

### 3.5. The effects of LLSOs compounds on EMT properties in different cancer types

We further sought to determine whether LLSOs can induce E-cadherin expression in cancer types beyond prostate cancer. MCF-7 and MDA-MB-231 for breast cancer, CALU-3 for lung cancer, HCT-116 for colon cancer and SKOV-3 and A2780 for ovarian cancer were used. To achieve this, we tested LLSO3 sensitivity and effect on E-cadherin expression in breast (MCF-7, MDA-MB-231), lung (CALU-3), colon (HCT116) and ovarian (SKOV-3, A2780) cancers.

Induction of E-cadherin by all three LLSOs was observed only in the MDA-MB-231 breast cancer cell line (**Figure 6A, B**), whereas in MCF-7 cells, only LLSO2 induced E-cadherin expression. No induction was detected in lung, colon, or ovarian cancer cell lines (CALU-3, HCT-116, SKOV-3, A2780; **Supplementary Figures 14** and 15).

**Figure 6.**
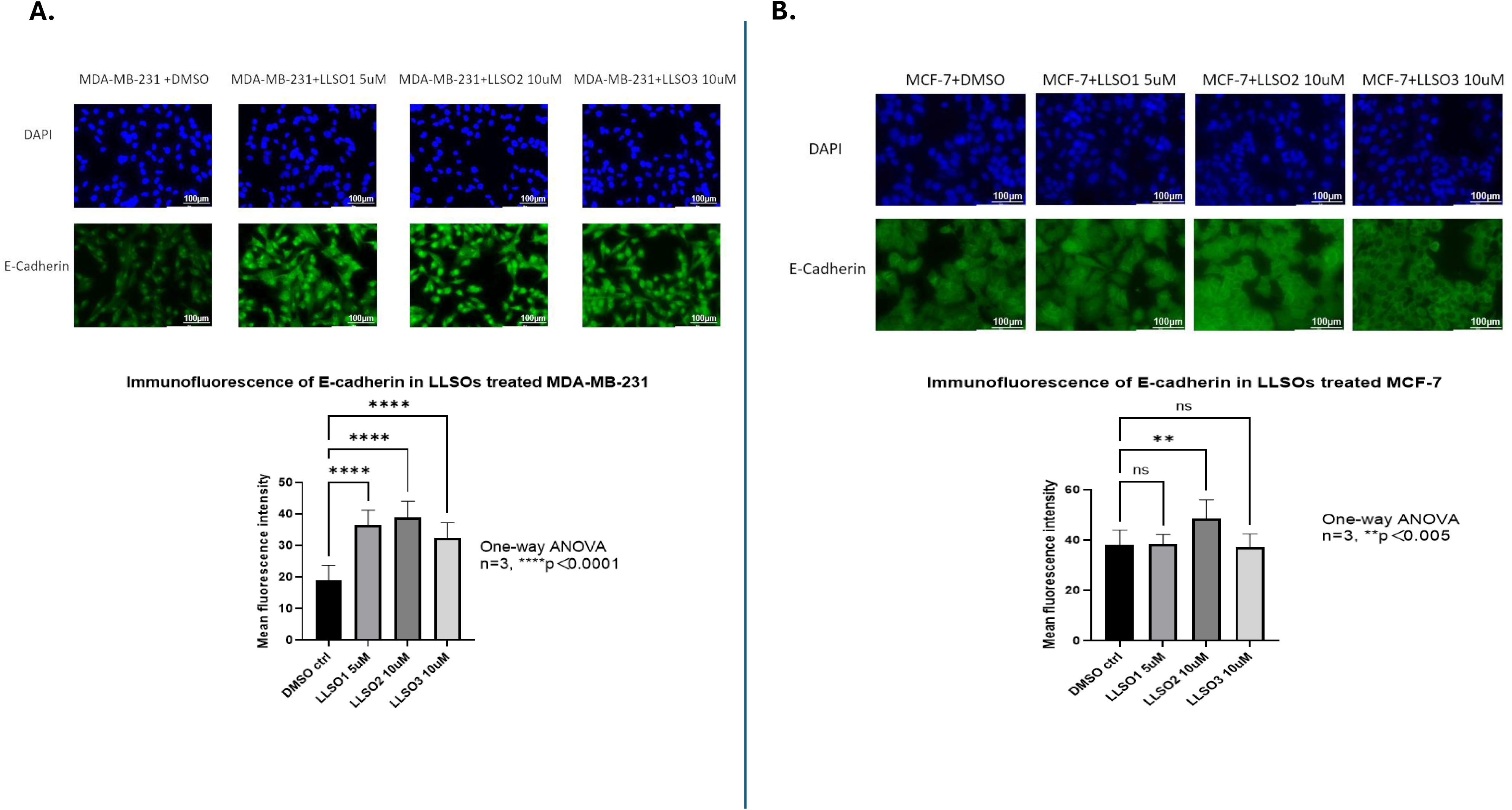
All LLSOs increased the expression of E-cadherin in MDA-MB-231 cells while only LLSO2 increased it in MCF-7. **A.** Immunofluorescence analysis for E-cadherin expression was performed following 48 hours treatment of MDA-MB-231 cells with DMSO, 5μM LLSO1, 10μM LLSO2, and 10μM LLSO3. MDA-MB-231 cells stained with mouse IgG were used as negative control and LNCaP cells were used as a positive control. All three LLSOs showed an increase in the E-cadherin expressing level. There are three wells repeated within each of the two independent experiments, n=3. Quantification of the E-cadherin signal was performed with ImageJ by using 9 fields of 3 repeat slides for each treatment. Photomicrographs were taken at 200x magnification; Scale bar is 100µm. **B.** Immunofluorescence analysis for E-cadherin expression was performed following 48 hours treatment of MCF-7 cells with DMSO, 5μM LLSO1, 10μM LLSO2, and 10μM LLSO3. MCF-7 cells stained with mouse IgG were used as negative control and LNCaP cells were used as a positive control. LLSO2 induced an increase in the expression level of E-cadherin, whereas LLSO1 and LLSO2 showed no effect on E-cadherin expressing level. There are three wells repeated within each of the two independent experiments, n=2. Quantification of the E-cadherin signal was performed with the ImageJ software by using 9 fields of 3 repeat slides for each treatment. Photomicrographs were taken at 200x magnification; Scale bar is 100µm.

### 3.6. Mechanism through which LLSOs compounds induce MET in PCa cells

In a previous paper, we have investigated the mechanism through which LLSO2 induces MET[10]. Here, we further characterised the mechanisms for LLSO1 and LLSO3.

#### LLSO1 signalling

As illustrated in the hypothetical LLSO1 signalling pathway depicted in **Supplementary Figure 16**, TTCC (T-type Calcium Channel) triggers signalling pathways through the elevation of intracellular Ca^2+^. After an influx of Ca^2+^, the calmodulin (CaM) and the downstream effector calmodulin kinase II (CaMKII), which are the proven transducers of TTCC activity, are activated[11]. Furthermore, studies have revealed that the Ras/Raf/MEK/ERK pathway is transiently activated after the activation of TTCC through the CaMKII, whereas inhibition of TTCC blocks the Akt/PKB pathway. Both signalling pathways were involved in the progression of DNA synthesis included in the cell cycle through controlling the cyclin/cyclin-dependent kinase (CDK) complexes[12–14]. Moreover, Wang et al also found that c-Src non- receptor tyrosine kinases function downstream of CaMKII in a signaling cascade and plays a critical role in many aspects of cancer progression including proliferation, migration and invasion in various cancer types[15, 16]. The CaMKII and CaMKIV activated by Ca2+/CaM were also found to be able to activate the cyclic AMP response element binding protein (CREB), overexpressed in diverse solid tumour types such as NSCLC, BrCc and glioblastoma, and have oncogenic potential by regulating a variety of cellular responses, including growth, proliferation, and survival[17]. Besides CREB, the transcription factor nuclear factor of activated T-cell (NFAT) could be activated by promoting activation of the well-known calcium and calmodulin dependent protein serine/threonine phosphatase calcineurin (CaN) through the influx of Ca^2+^ [18, 19].

The mechanism by which LLSO1 signals to increase the expression level of E-cadherin was determined by using the immunofluorescence described above. CaM as calcium-sensing protein has more than 200 distinct substrates, and the essential targets of CaM-dependent pathways required for cell proliferation remain largely elusive[20, 21]. Hence, one of the key regulators of intracellular calcium signaling, CaN, was first targeted as activation of CaN has been considered to play a pro-tumorigenic role, which is linked to increased cells activity, including promoting proliferation, migration, and invasion in diverse cancers like BrCc, PCa, LUAD and COAD[22].

The widely used CaN inhibitor, FK506 (Tacrolimus), has been shown to suppress the proliferation and migration of several cancers including PCa and BrCc[23–25]. We therefore treated PC3 cells with FK506, alone and in combination with LLSO1, to assess whether inhibition of CaN recapitulates the effects of LLSO1 on E-cadherin expression. As shown in **Figure 7A–B**, FK506 treatment—whether alone or with LLSO1—led to increased E-cadherin expression, comparable to that observed with LLSO1 alone. These findings suggest that LLSO1 may exert its effects through the CaN signaling pathway.

**Figure 7.**
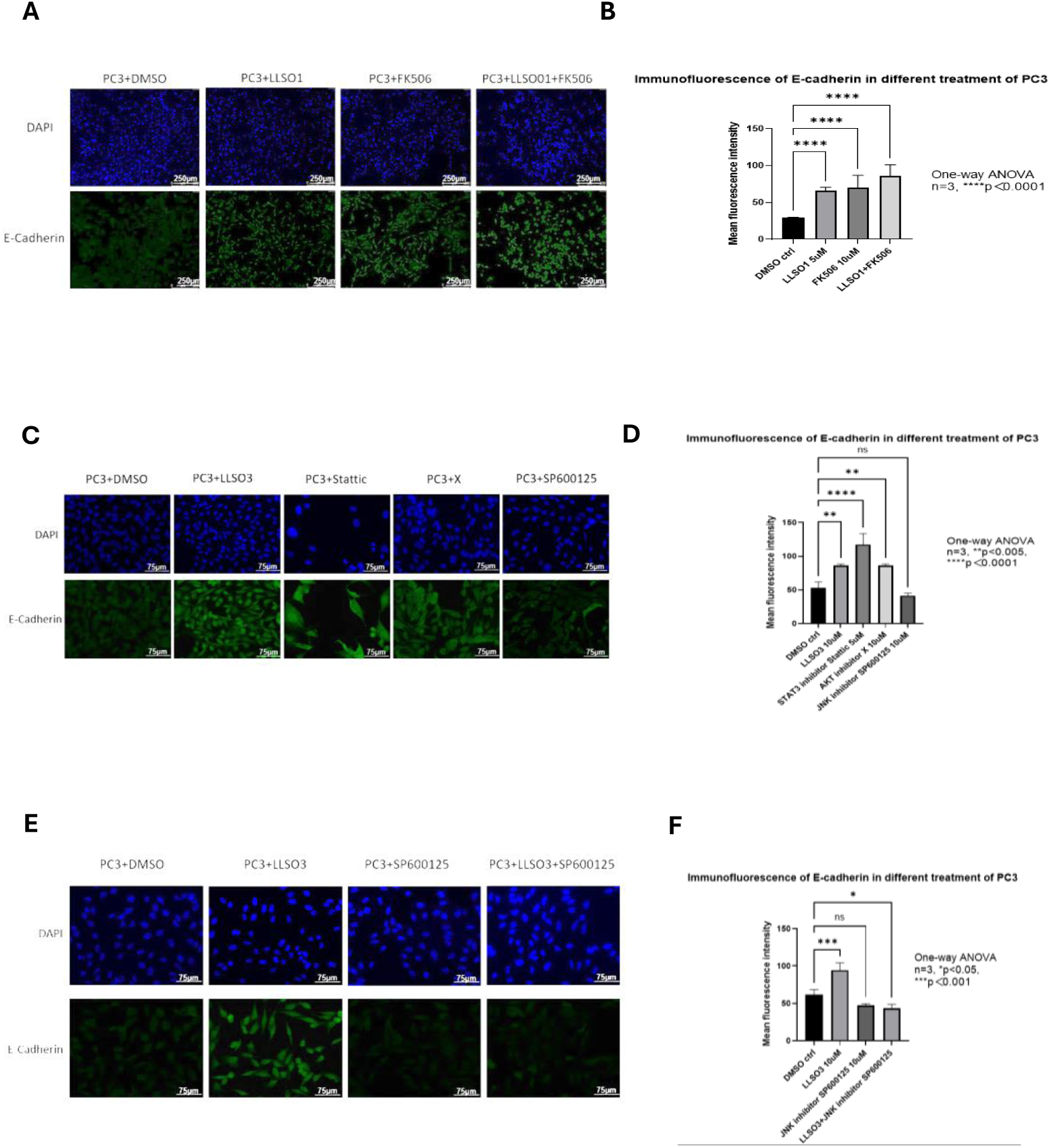
Signalling pathways used by LLSO1 and 3 in PC-3 cells. ***Left panel*: Inhibition of CaN increased E-cadherin expression in PC3 cells.** **A.** Images of PC3 cells treated separately with 10μM of LLSO1, 40μM of FK506, and the combination of LLSO1 and FK506. All images were taken at 100x magnification. The scale bar is 250µm. Representative examples from at least 9 microscopic fields per experiment, experiments repeated 3 times with triplicate per treatment. **B.** Quantification of immunofluorescence for E-cadherin expression was performed after 48 hours of treatments of PC3 cells with DMSO, 10μM of LLSO1, 20μM of FK506, or the combination of LLSO1 and FK506. All these treatments showed an increase in the level of E-cadherin expression. n=3, data was analysed by one-way ANOVA. ***Middle panel*: Inhibition of STAT3 increased E-cadherin expression in PC3 cells, but not inhibition of AKT or JNK.** **C.** Images of PC3 cells treated separately with 10μM of LLSO3, 5μM of STAT3 inhibitor Stattic, 10μM of AKT inhibitor X, and 10μM of JNK inhibitor SP600125. All images were taken at 200x magnification. The scale bar is 250µm. Representative examples from at least 9 microscopic fields per experiment, experiments repeated 3 times with triplicate per treatment. **D.** Quantification of immunofluorescence for E-cadherin expression was performed after 48 hours treatments of PC3 cells with DMSO, 10μM of LLSO3, 5μM of STAT3 inhibitor Stattic, 10μM of AKT inhibitor X, and 10μM of JNK inhibitor SP600125. All these treatments showed an increase on E-cadherin expression level. n=3, data was analysed by one-way ANOVA. ***Right panel*: LLSO3 uses JNK signalling pathway to increase E-cadherin expression in PC3 cells.** **E.** Images of PC3 cells treated separately with 10μM of LLSO3, 10μM of AKT inhibitor X, the combination of LLSO3 and X, 10μM of JNK inhibitor SP600125, and the combination of LLSO3 and SP600125. All images are taken at 200x magnification. The scale bar is 250µm. Representative examples from at least 9 microscopic fields per experiment, experiments repeated 3 times with triplicate per treatment. **F.** Quantification of immunofluorescence for E-cadherin expression was performed after 48 hours treatments of PC3 cells with DMSO, 10μM of LLSO3, 10μM of AKT inhibitor X, the combination of LLSO3 and X, 10μM of JNK inhibitor SP600125, and the combination of LLSO3 and SP600125. All these treatments showed an increase on E-cadherin expression levels except for the treatment of 10μM of JNK inhibitor SP600125, and the combination of LLSO3 and SP600125. n=3, data was analysed by one-way ANOVA.

#### LLSO3 signalling

An increasing number of studies have described that several opioid receptors and receptor subtypes are widely implicated in various types of cancer cells and are associated with cancer aggressiveness, particularly G-protein-coupled receptors including mu-opioid receptor (MOR), kappa-opioid receptor (KOR), and delta-opioid receptor (DOR)[26, 27]. Based on the illustrated hypothetical signalling pathway depicted in **Supplementary Figure 17**, it is possible that LLSO3, functioning as a non-selective opioid receptor antagonist, possesses the potential to affect all these pathways. Since STAT3 and AKT both have the ability to regulate certain types of cancer progression through modulating EMT, and are both present in all three opioid receptor pathways, whether the STAT3 and AKT are involved in the mechanisms of LLSO3 inhibiting PCa cells growth and migration, and EMT regulation were firstly assessed[28, 29]. In addition, a growing body of studies have indicated that activation of JNK play a critical role in inducing EMT and promote migration and invasion of several cancer cells, including PCa, BrCc, and COAD[30–32]. Therefore, whether STAT3, AKT and JNK are involved in the effect of LLSO3 on EMT regulation were also investigated by using Stattic as STAT3 inhibitor, X as AKT inhibitor, and SP600125 as JNK inhibitor.

As shown in **Figure 7C-D**, treatment of PC3 cells with both Stattic and LLSO3 increased E-cadherin expression above the level of LLSO3 alone, indicating that STAT3 inhibition may play a role in mediating LLSO3’s effect. In contrast, treatment of PC3 cells with both LLSO3 and AKT had no effect on E-cadherin expression, whilst co-treatment with LLSO3 and JNK in fact prevented prevented the upregulation of E-cadherin induced by LLSO3 (**Figure 7E, 7F**). These findings suggest that LLSO3 likely exerts its EMT-inhibitory effects via the JNK signalling pathway rather than the AKT pathway.

## 4. Discussion

### 4.1 The applications and limitations of cancer therapies targeting EMT

EMT plays a significant role in cancer progression that includes metastasis and invasion. EMT is also critical in drug resistance acquisition in cancer treatment and in immunosuppression and refractory responses to chemotherapy. Consequently, EMT has become a highly valued target for the development of novel cancer therapeutic strategies. EMT-targeting therapeutic strategies may be able to prevent cancer cell invasion and dissemination, as well as reduce cancer stemness and increase the efficacy of traditional chemotherapeutics[33].

Common EMT-targeting strategies including:

a. *Blocking key signalling pathways*

There are several key signalling pathways involved in the regulation of EMT, which include TGF-β, RTK, EGF, FGF, IGF, PDGF, Wnt, Notch, Hedgehog, MAPK/ERK, Hippo, matrix signalling pathway and hypoxia signalling pathway[34–37]. These signalling pathways regulate EMT by modulating several transcription factors. For example, Luteolin reverse EMT through inhibiting the Notch signalling pathway, consequently suppressing the progression of gastric cancer[38].

*b. Targeting the EMT-inducing transcription factors (EMT-TFs)*

Transcription factors that bind to the genes encoding adherens junction and tight junction molecules, such as E-cadherin, claudins, and occluding, are indispensable drivers of EMT. EMT-TFs such as the Snail1/2, ZEB1/2, Twist1/2 and LEF-1 have been identified as master regulators of EMT through directly repressing E- cadherin[34]. In addition, EMT-TFs without directly targeting CDH1 gene promoter suppression including Goosecoid, FOXC2, TCF4[39–41] and many other transcriptional repressors are also inducing EMT. Although inhibiting some EMT-TFs may block or reverse EMT, it may also risks disrupting anti-apoptotic signalling and other normal stem cell regulation[42].

*c. Regulating EMT markers*

A growing body of research indicates that directly targeting the epithelial cells by enhancing the functions of epithelial-specific proteins such as E-cadherin and cytokeratin can repress EMT and boost MET features, thereby reducing the tumour cells invasion and metastasis[43]. Furthermore, targeting the mesenchymal cells directly through inhibiting the expression of mesenchymal markers such as N- cadherin, vimentin, and fibronectin, which have also been shown the suppression of EMT[44–46].

For instance, agents like VNLG-152, a novel retinamide could inhibit EMT in PCa by reducing the expression of N-cadherin, β-catenin, vimentin, claudin and Twist while simultaneously increasing E-cadherin expression[47]. However, reversion to epithelial phenotype may stimulate the formation of secondary metastases from the primary tumour cells that have already disseminated[33].

*d. Modulating microRNAs*

MicroRNAs (miRs) are well established as crucial regulators of EMT in epithelial cells that target multiple EMT/MET components[48]. Due to miRNA can directly bind and suppress EMT-TFs, EMT signalling pathways, and key EMT marker proteins[49], thus miRNAs can either inhibit EMT or promote MET. The miR-200 family is a well-studied pathway of miR-mediated EMT suppression that works primarily through directly targeting the transcriptional repressors ZEB1/2[50, 51].

### 4.2 Potential signalling pathway involved in the LLSO1 effects on PCa cells

The immunofluorescence results showed that LLSO1 increased E-cadherin expression in PC3 cells (Figure 7A), particularly in combination with the CaN inhibitor, suggesting involvement of the CaN pathway. Even though the distribution of CaN has been widely found in various mammalian tissues, activation of CaN and its downstream targets have been implicated in a variety types of cancer progression including colon, breast, lung and hepatocellular carcinoma[52–55]. Notably, the pro- tumorigenesis role for CaN signalling in cancer have focused principally on NFAT, which is activated by CaN to promote cells proliferation[18, 56, 57]. Since CaN dephosphorylates cytoplasmic NFAT to activate NFAT translocation into the nucleus and consequently activates the transcription of downstream gene targets including Src to promote EMT progression[56, 58], inhibition of CaN supress EMT progression via NFAT might be a potential target. However, Dantal et al discovered that the incidence of cancer in patients has significant increased after immunosuppressant treatment[59], and CaN inhibitor like FK506 has been widely used immunosuppressant clinically. CaN inhibition has shown contradictory roles, promoting EMT in some cases[60] while inhibiting it in others[61] . Additionally, it has also been found FK506 might inhibit the growth of PCa, BrCc and bladder cancer[62–64].

In summary, our finding suggest LLSO1 suppresses EMT is highly likely through CaN/NFAT.

### 4.3 Potential signalling pathways involved in the LLSO3 effects on PCa cells

As shown in Supplementary Figure 16, LLSO3 could behave as a non-selective opioid receptor antagonist, which have the potential of inhibiting several opioid receptors including mu-opioid receptor (MOR), kappa-opioid receptor (KOR), and delta-opioid receptor (DOR)[26, 27].

Immunofluorescence results shown that LLSO3 upregulated E-cadherin (Figure 2A), which indicated the inhibition of either MOR, KOR or DOR are responsible for the activation of E-cadherin expression in PC3 cells. The upregulation of E-cadherin was also present when STAT3 signalling was inhibited (Figure 7C-D), which suggests that the mechanism of LLSO3 to inhibit PCa cellular’ growth and migration, and upregulate E-cadherin might occur through inhibiting the STAT3 signalling. This has been previously investigated by Cho et al. and Hu et al., who showed that suppression of STAT3 in Pca cells inhibited TGFβ-induced EMT and Pca cells invasion[65, 66]. Therefore, the mechanism of LLSO3 inducing E-cadherin expression and inhibiting PC3 cells involving the STAT3 signalling is plausible.

On the other hand, while AKT and JNK inhibitors alone did not affect E- cadherin, JNK inhibition abrogated LLSO3’s effect (Figure 7E-F), implying that JNK signalling might modulate EMT in a context-dependent manner. The mechanism of LLSO3 to upregulated E-cadherin via JNK pathway seems conceivable, as multiple studies have found that JNK signalling is implicated in EMT regulation in various types of cancer, including BrCc, COAD, Pca and cervical cancer[30, 31, 67, 68].

To further investigate the mechanism of LLSO3 inhibition of growth and migration of PC3 cells and EMT regulation, either upstream or downstream mediators of STAT3 such as JAK1/2 and PI3K, as well as for JNK like GSK3β, p53 and β-catenin, can be disrupted either by pharmacological inhibitors or genetic silencing. However, whether STAT3 or JNK are involved in the inhibitory effect of LLSO3 on migration and cellular growth of DU145 cells and E-cadherin upregulation has yet to be determined.

### 4.4 The association between LLSO3, EMT properties, and tumour growth

The low-dose naltrexone is used as a treatment for pain perception and in the inhibition of the opioid effect on the central nervous system. However, only a few research studies have implicated naltrexone in cancer therapy and none of them have associated this drug as a possible treatment for Pca, even if Zagon I.S. and McLaughlin P.J. reported the anti-tumour growth effect of Naltrexone long time ago[69]. The use of opioid receptor antagonists may be a potential opportunity for treatments aimed at inhibiting cancer progression, and the mechanisms by which LLSO3 inhibits the growth and migration of PC3 might be similar to the canonical mechanisms for targeting opioid receptors.

Furthermore, Lennon FE et al. described that the Mu-opioid receptor promotes opioid and growth factor-induced EGF receptor signalling, including PI3K, Akt and STAT3, and consequently stimulates cancer cell growth, proliferation, metastasis and EMT in lung cancer progression[70]. In addition, Tripolt, S et al. discovered that the expression of delta-opioid receptor is not only implicated in poor prognosis for breast cancer patients, but also promoted the metastasis and migration of BrCc through activation of oncogenic JAK1/2-STAT3 signalling and promoting EMT[71]. Taking all these studies into account, it may be that the expression of opioid receptors promotes cancer progression while triggering the progression of EMT, which might explain why LLSO3 reduces Pca tumour xenografts growth through affecting EMT, as the naltrexone hydrochloride is a non-selective opioid receptors antagonist. LLSO3 not only inhibits the growth, proliferation and migration of Pca cells *in vitro*, but also suppresses Pca tumour growth and affects EMT *in vivo*; LLSO3 might be a potential therapeutic drug for Pca through blocking opioid receptors and inhibiting the process of EMT.

In conclusion, our evidence warrants further investigation of LLSO compounds in the development of tumour therapeutics. This might include the repurposing of the compounds themselves or further investigations into the signaling pathways they are involved to discover common denominators and key regulators that could further be used in drug development.

## Acknowledgments

We wish to thank Megan Hill and Tattie Barclay-White for assistance with experiments involving LLSO1.

---

**Supplementary Figure 1.**
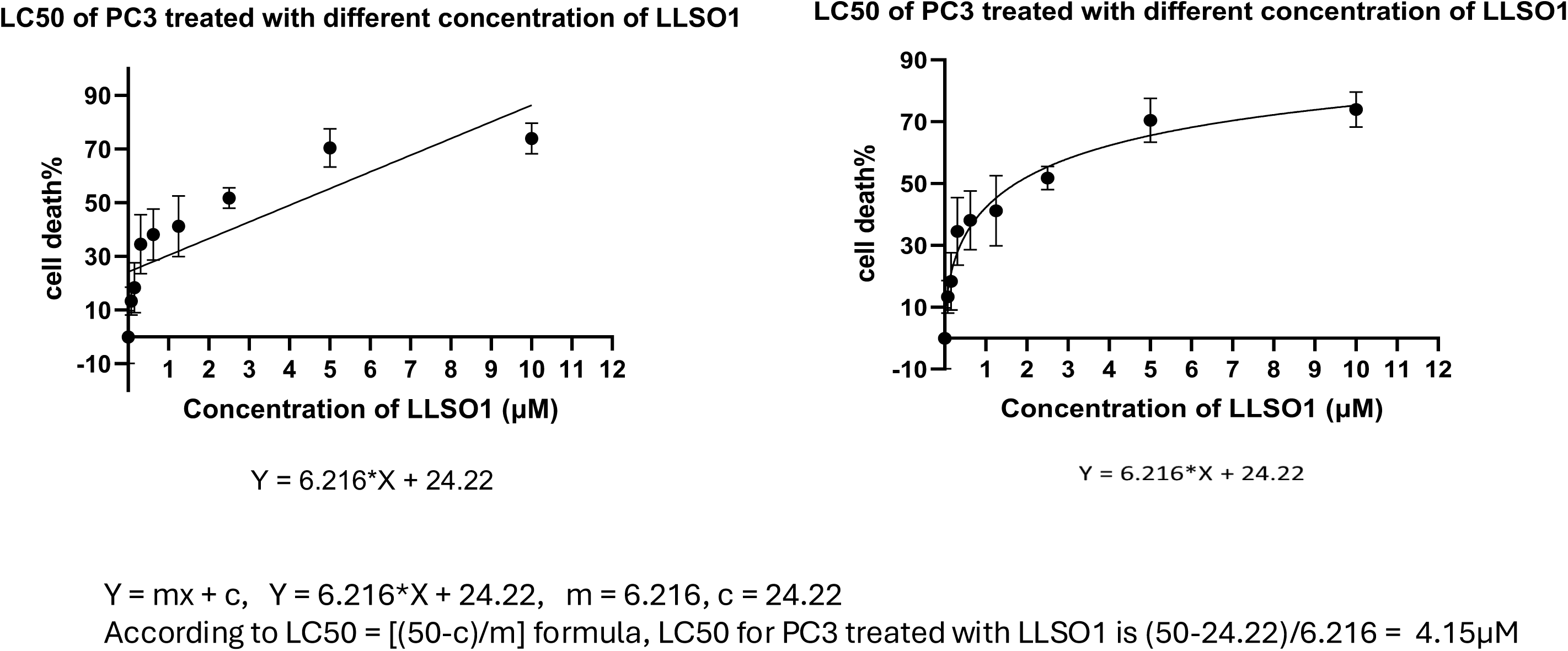
The LC50 for PC3 treated with different concentration of LLSO1 is 4.15 μM. MTT assay was performed on PC3 cells following 48 hours of treatment with DMSO or LLSO1 at concentration of 0.078125μM, 0.15625μM, 0.3125μM, 0.625μM, 1.25μM, 2.5μM, 5μM, and 10μM. Cell viability was assessed by measuring absorbance 590nm. Data represent mean ± SD of six replicates per condition. **Left panel:** The linear regression model described by the equation Y = 6.216*X + 24.22, was used to calculate LC50 via the fomula LC50 = (50-c)/m, yielding a value of 4.15μM. **Right panel:** the non-linear regression mdel more accurately reflects the sigmoidal dose-response relationship observed in cytotoxicity assays.

**Supplementary Figure 2.**
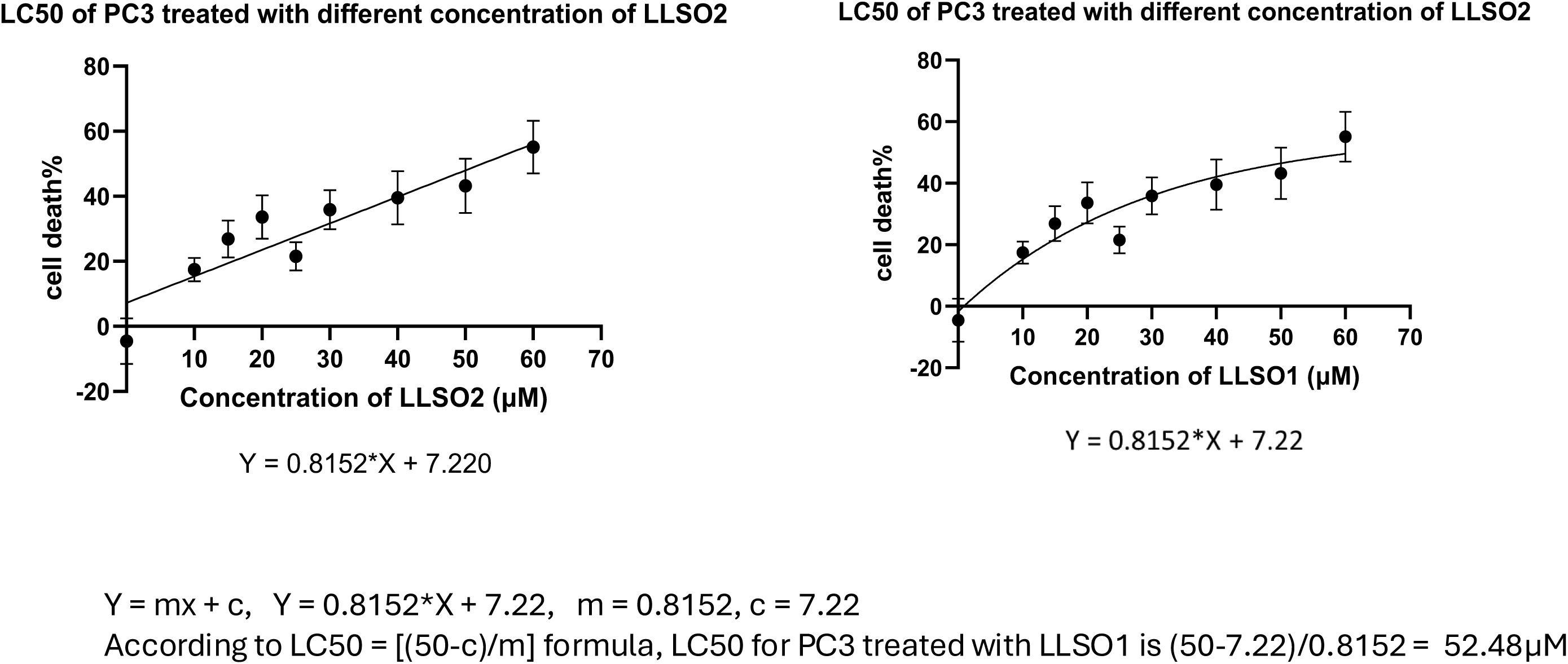
The LC50 for PC3 treated with different concentration of LLSO2 is 52.48 μM. MTT assay was performed performed on PC3 cells following 48 hours of treatment with DMSO or LLSO2 at concentration of 10μM, 15μM, 20μM, 25μM, 30μM, 40μM, 50μM, and 60μM, and viability was assessed at OD = 590nm. Data represent mean ± SD (n = 6 replicates per condition). **Left panel:** LC50 was calculated using the linear regression model (Y = 0.8152*X + 7.22; LC50 = (50 − c)/m), yielding a value of 52.48μM. **Right panel:** the non-linear regression mdel more accurately reflects the sigmoidal dose-response relationship observed in cytotoxicity assays.

**Supplementary Figure 3.**
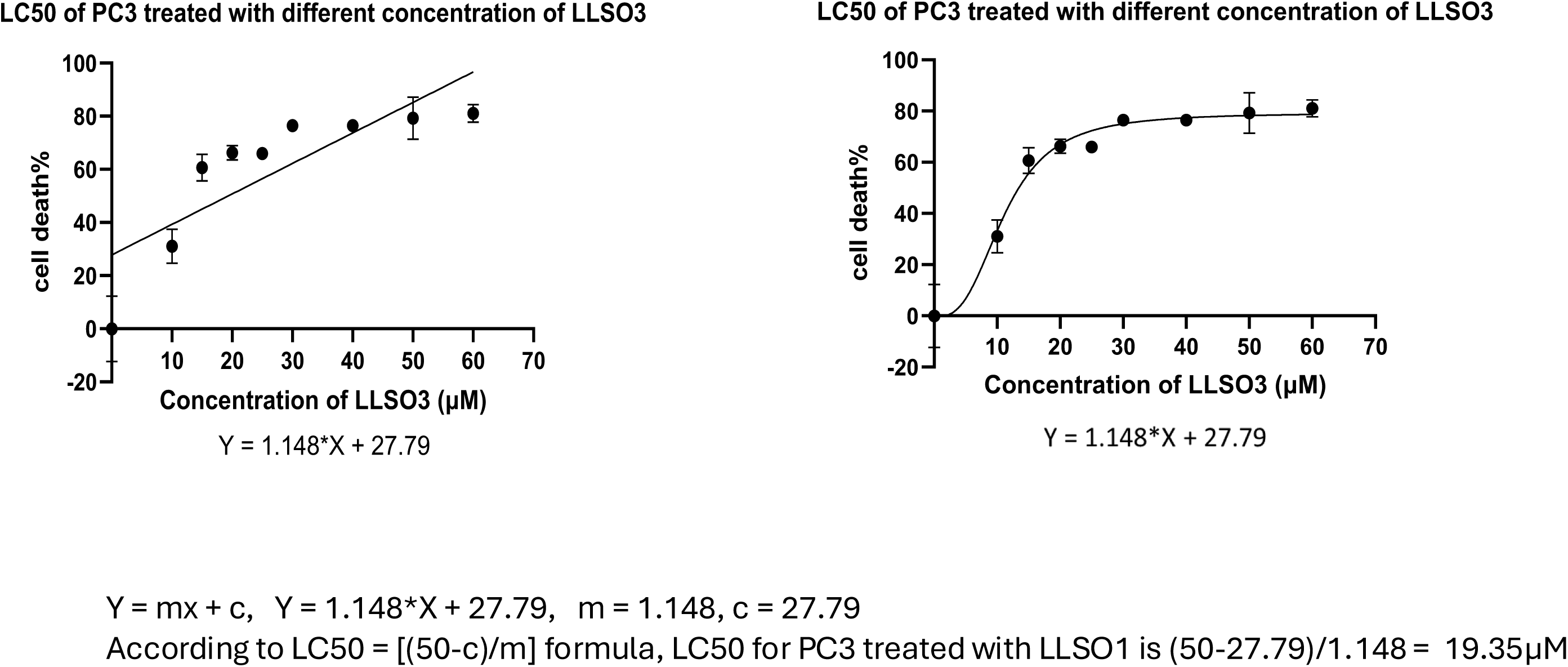
The LC50 for PC3 treated with different concentration of LLSO3 is 19.35 μM. . MTT assay was performed performed on PC3 cells following 48 hours of treatment with DMSO or LLSO3 at concentration of 10μM, 15μM, 20μM, 25μM, 30μM, 40μM, 50μM, and 60μM, and viability was assessed at OD = 590nm. Data represent mean ± SD (n = 6 replicates per condition). **Left panel:** LC50 was calculated using the linear regression model (Y = 1.148*X + 27.79; LC50 = (50 − c)/m), yielding a value of 19.35μM. **Right panel:** the non-linear regression mdel more accurately reflects the sigmoidal dose-response relationship observed in cytotoxicity assays.

**Supplementary Figure 4.**
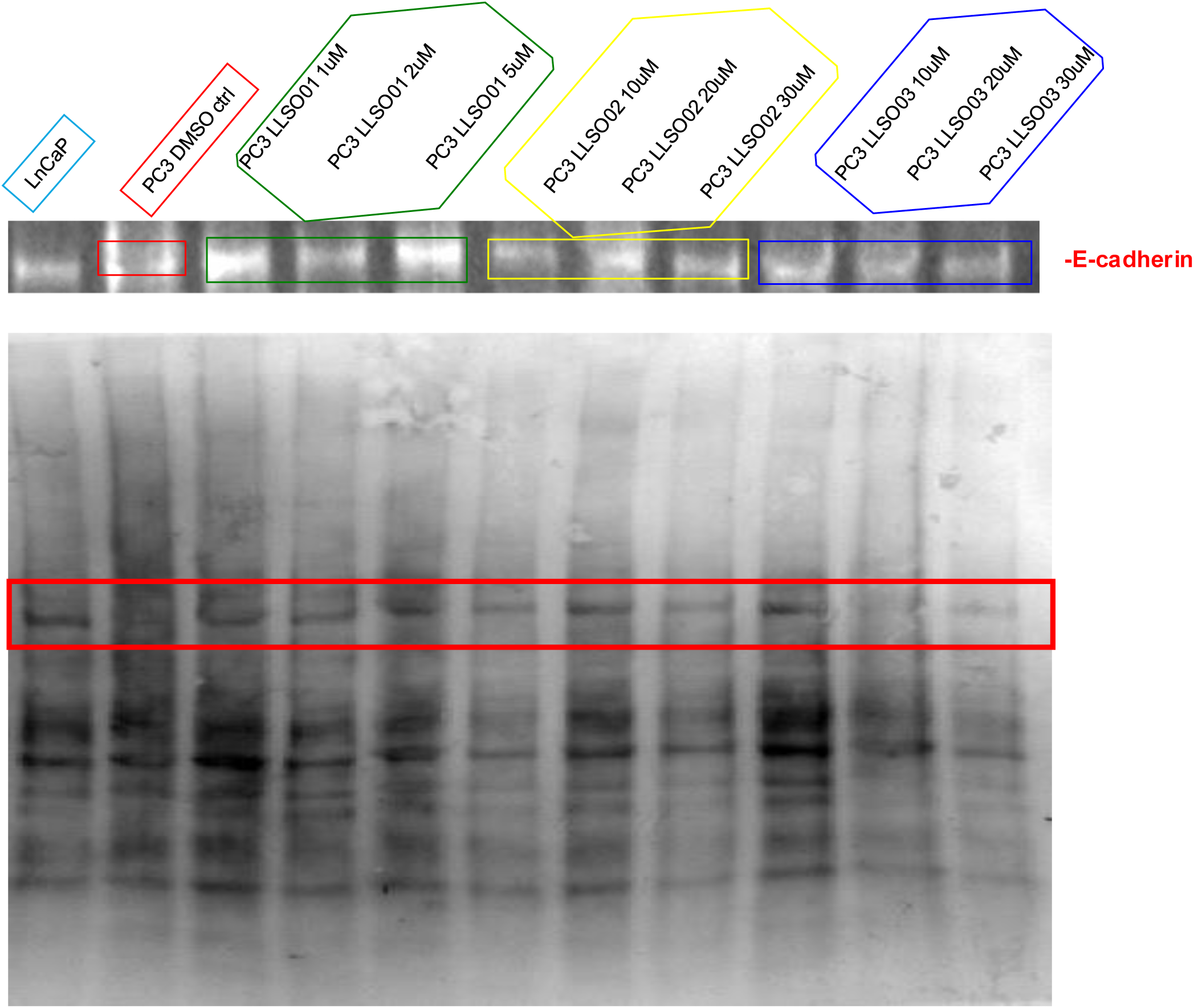
LLSOs increase the expression of E-Cadherin in PC3 cells. Western blot for E-Cadherin expression was performed with proteins extracted following 48 hours treatments with DMSO and different concentrations of LLSOs in PC3 cells. Upper panel is the antibody signal; lower panel is the total protein quantification on the transfer membrane using BioRad’s GelDoc EZ imaging device.

**Supplementary Figure 5.**
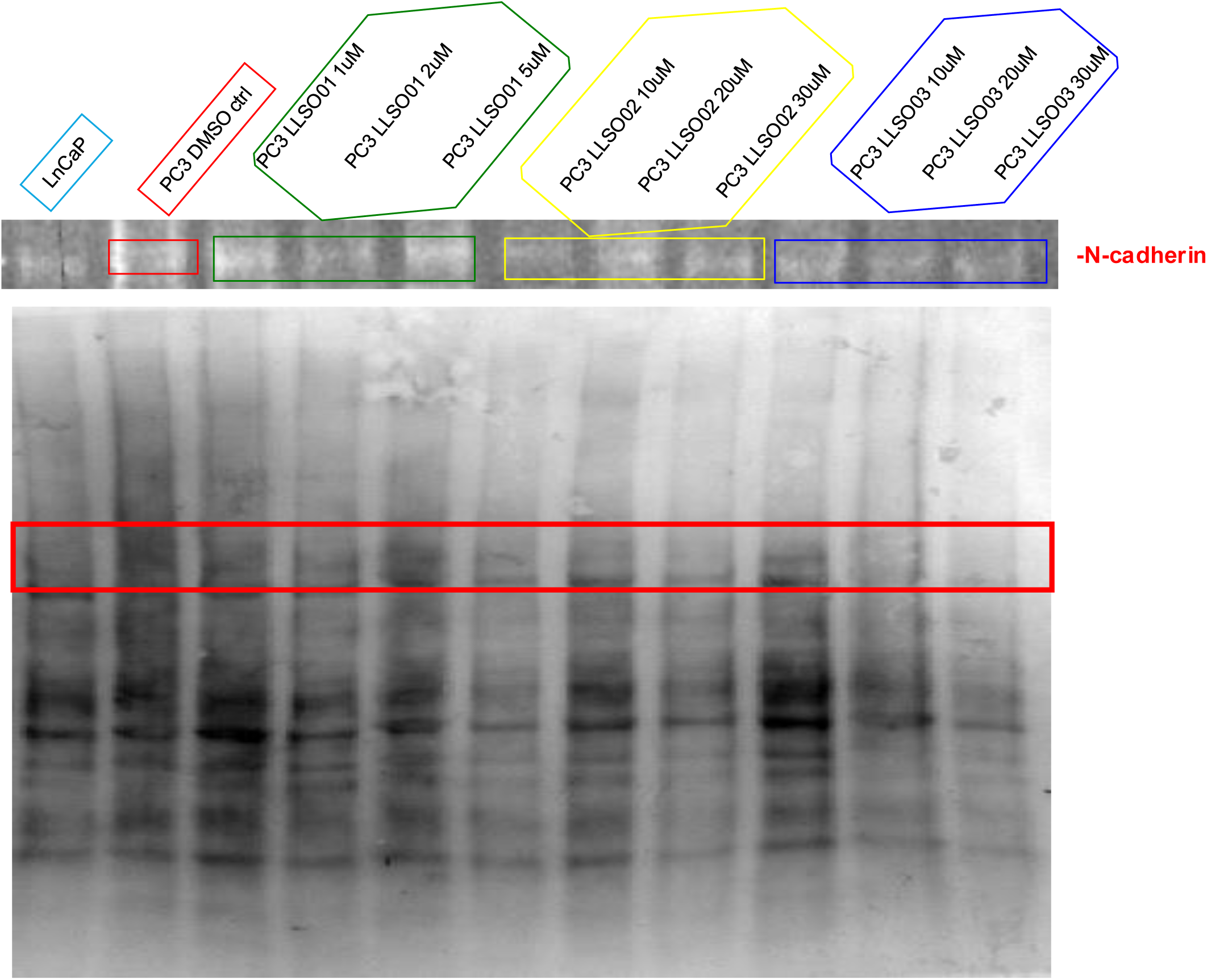
LLSOs decreases the expression of N-Cadherin in PC3 cells. Western blot for N-Cadherin expression was performed with proteins extracted following 48 hours treatments with DMSO and different concentrations of LLSOs in PC3 cells. Upper panel is the antibody signal; lower panel is the total protein quantification on the transfer membrane using BioRad’s GelDoc EZ imaging device.

**Supplementary Figure 6.**
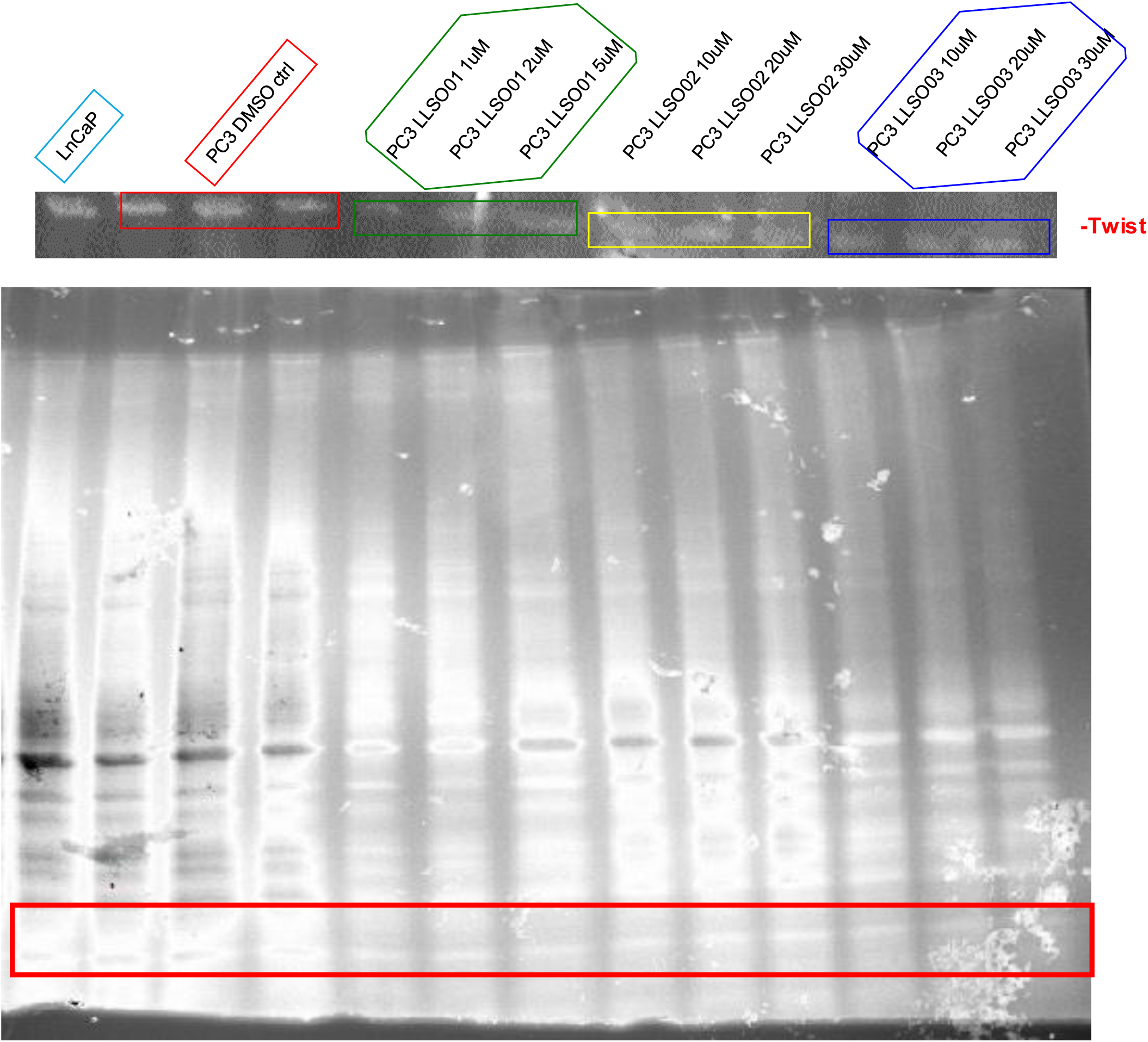
LLSOs decreases the expression of Twist in PC3 cells. Western blot for Twist expression was performed with proteins extracted following 48 hours treatments with DMSO and different concentrations of LLSOs in PC3 cells. Upper panel is the antibody signal; lower panel is the total protein quantification on the transfer membrane using BioRad’s GelDoc EZ imaging device.

**Supplementary Figure 7.**
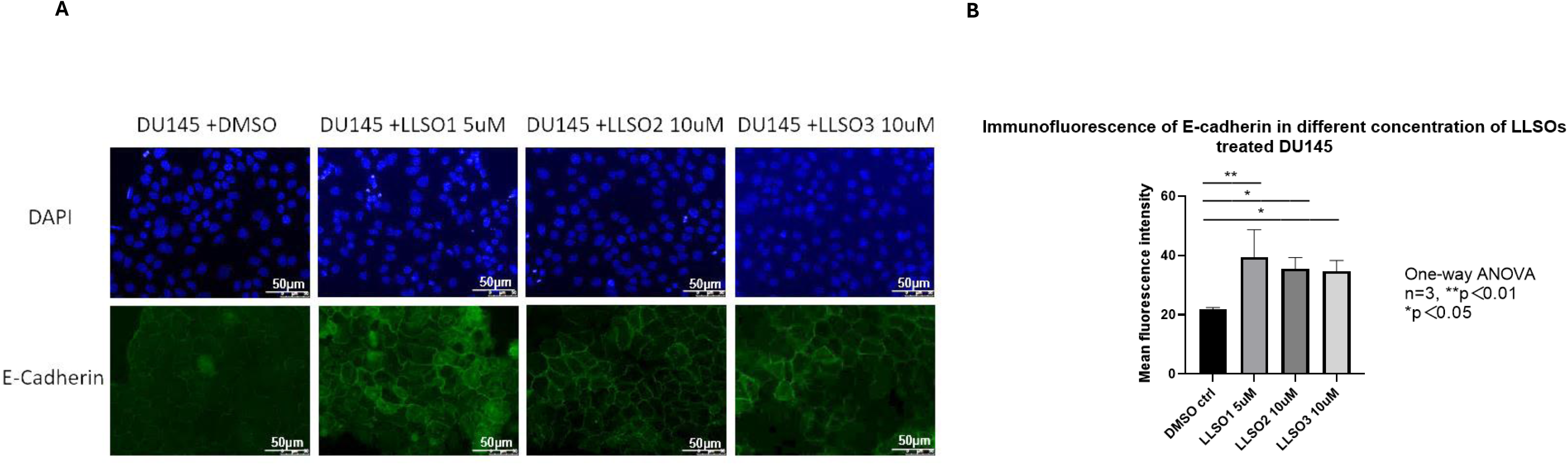
Different concentration of LLSOs slightly increase the expression of E-cadherin in DU145 cells. Immunofluorescence was performed following 48 hours pre-treatment of DMSO and 5uM of LLSO01, 10uM of LLSO02 and 10uM of LLSO03 in DU145 cells. **A.** Examples of microscopic fields taken at 20x magnification **B.** Normalized E-cadherin expression of LLSO01 5µM, LLSO02 10µM, LLSO03 10µM treated DU145 cells and DMSO (as control) treated DU145 cells. n=3, data was analysed by one-way ANOVA

**Supplementary Figure 8.**
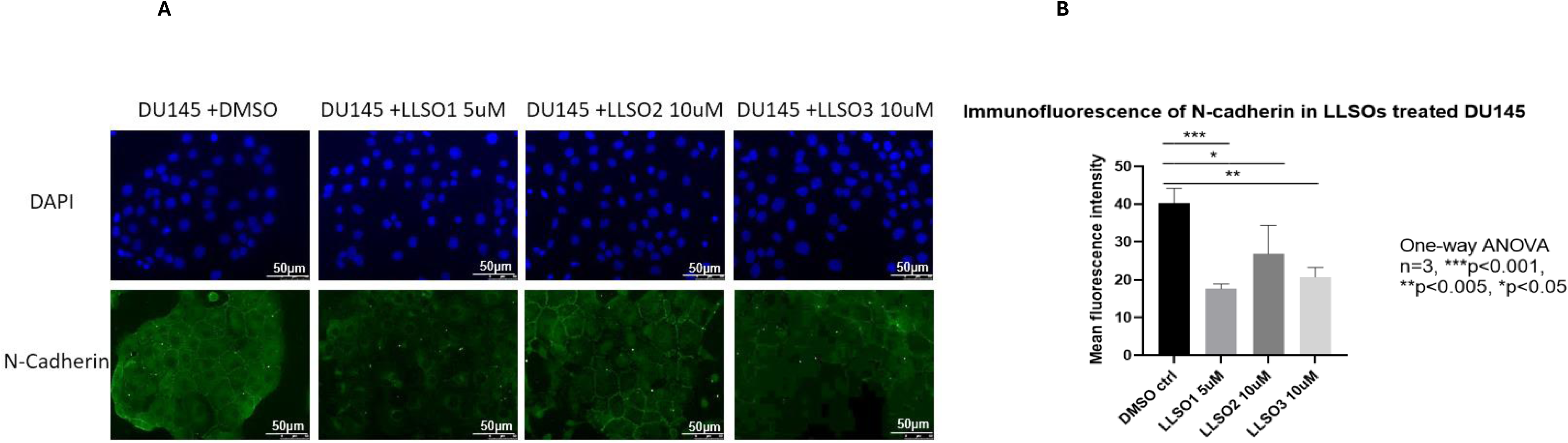
Different concentration of LLSOs decrease the expression of N-cadherin in DU145 cells. Immunofluorescence was performed following 48 hours pre-treatment of DMSO and 5uM of LLSO01, 10uM of LLSO02 and 10uM of LLSO03 in DU145 cells. **A.** Examples of microscopic fields taken at 20x magnification **B.** Normalized N-cadherin expression of LLSO01 5µM, LLSO02 10µM, LLSO03 10µM treated PC3 cells and DMSO (as control) treated DU145 cells. n=3, data was analysed by one-way ANOVA

**Supplementary Figure 9.**
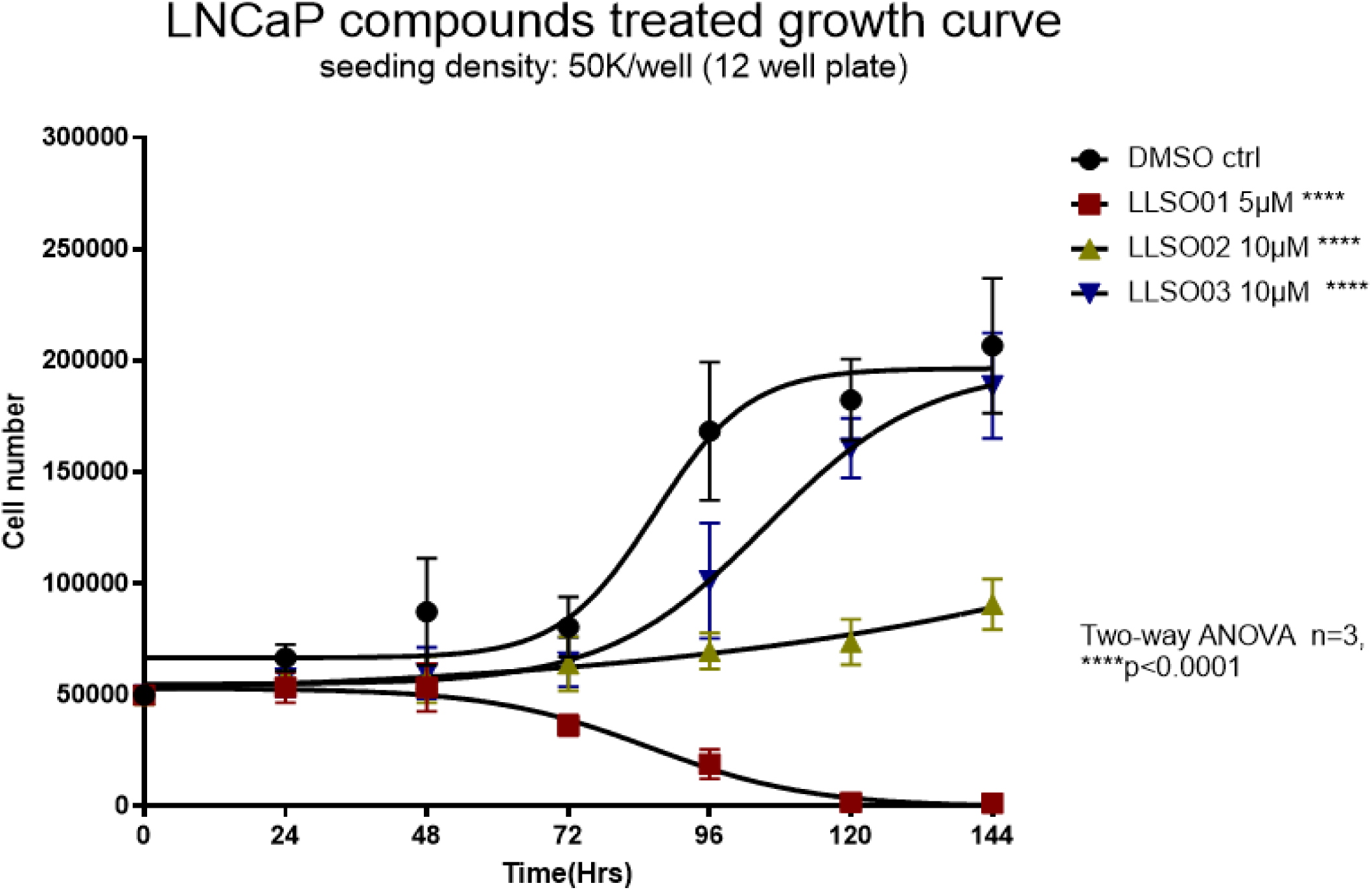
LLSOs inhibit LNCaP cells’ growth. Growth curve of LLSOs and DMSO treated LNCaP cells. For the three treatments – LLSO1 is toxic to LNCaP cells, LLSO2 dramatically inhibits LNCaP cell growth, and LLSO3 slightly slows the growth rate. The medium with LLSOs treatment was refreshed every 48 hours after 24 hours of seeding LNCaP cells, and cells were counted every 24 hours after seeding in the plate. Three individual experiments with three repeats for each treatment in each individual experiment, data were analyzed by two-way ANOVA. n=9, ****p<0.0001

**Supplementary Figure 10.**
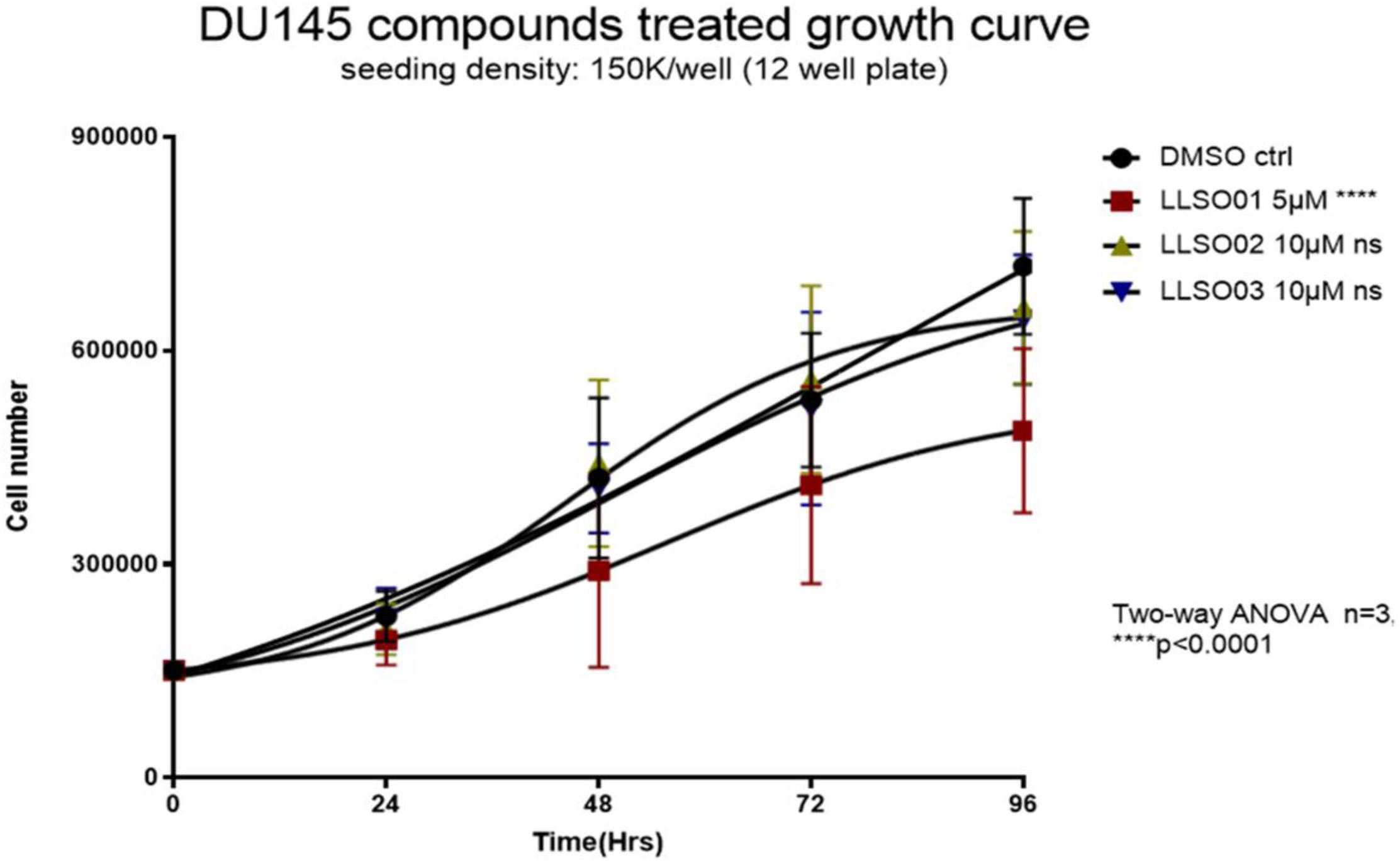
LLSOs affect DU145 cells’ growth. Growth curve of LLSOs and DMSO treated DU145 cells. LLSO1 slightly inhibits the growth of DU145 cells, whereas both LLSO2 and LLSO3 show no significant effect on cell growth. Medium with LLSOs treatment was refreshed every 48 hours after 24 hours of seeding DU145 cells, and cells were counted every 24 hours after seeding in the plate. Three individual experiments with three repeats for each treatment in each individual experiment, the data were analysed by two-way ANOVA. n=9, ****p<0.0001, ns, not significant.

**Supplementary Figure 11.**
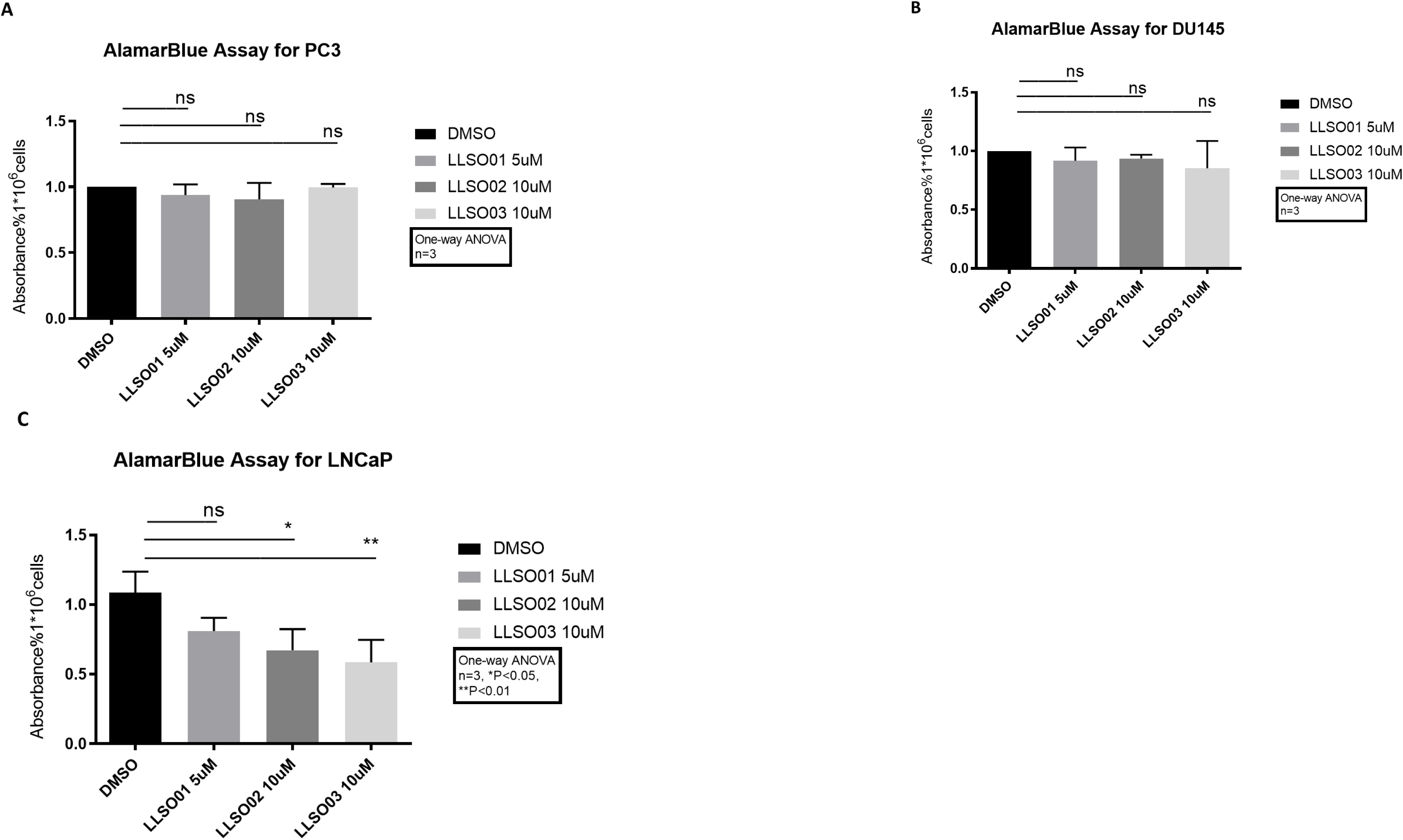
LLSOs slightly reduce cell viability in LNCaP cells, while no effect is observed on both PC3 and DU145 cell viability. AlamarBlue assay was performed following 48 hours of pre-treatment of DMSO and LLSOs in PC3 cells, DU145 cells and LNCaP cells. Absorbance rate on AlamarBlue assay of (A) PC3, (B) DU145 and (C) LNCaP treated with 5µM LLSO1, LLSO2 10µM, LLSO3 10µM and DMSO (as control). n=3, twelve replicated within each experiment, data were analysed by one-way ANOVA. ns, not significant.

**Supplementary Figure 12.**
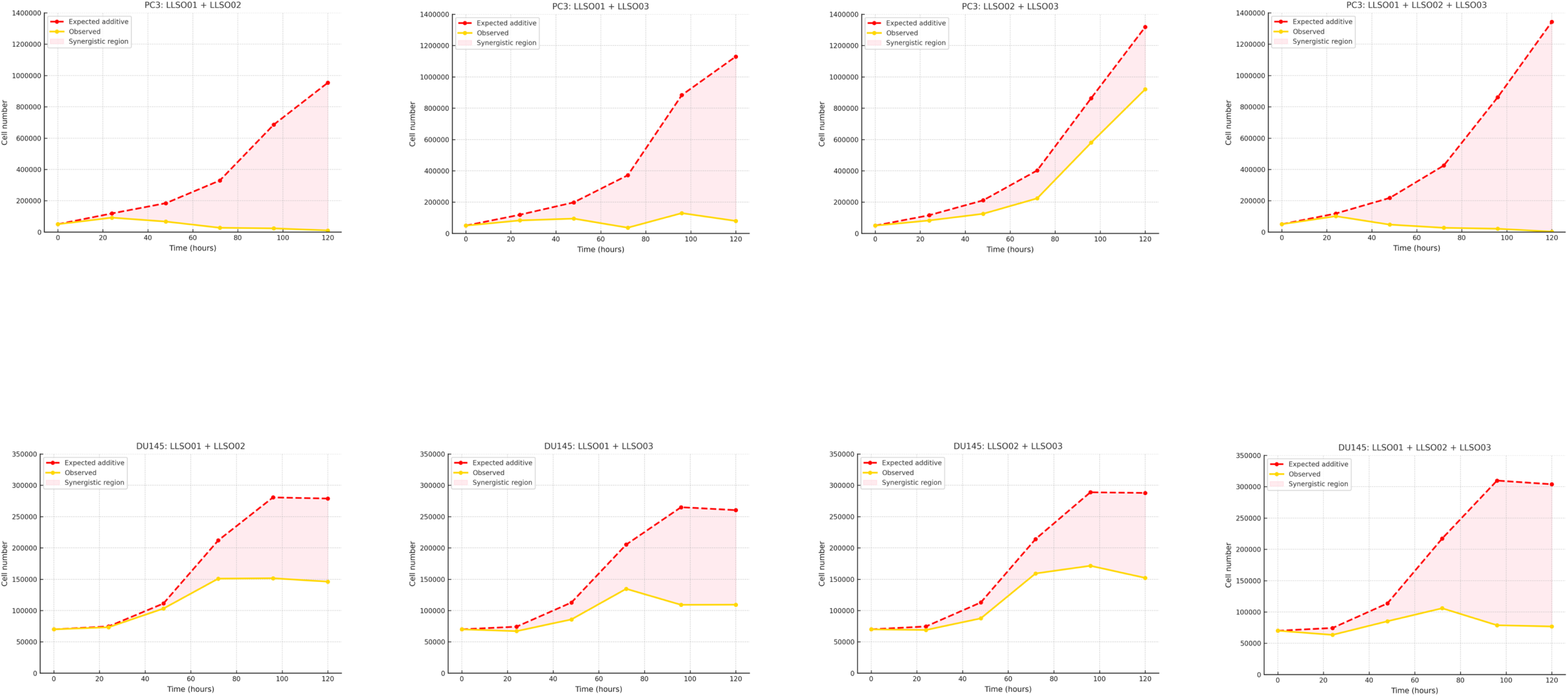
Bliss independence analysis in PC3 (upper panel) and DU145 (lower panel) cells.

**Supplementary Figure 13.**
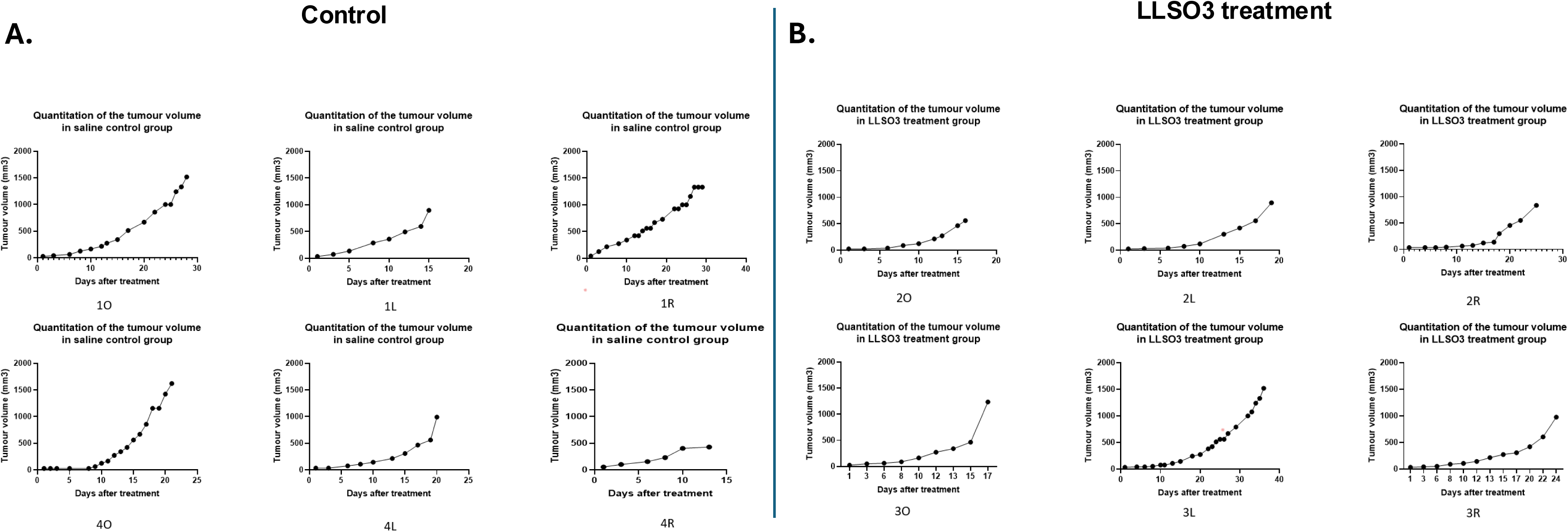
Growth curves monitoring the tumour volume in saline control group (A) and LLSO3 treatment group (B) Tumour volumes were estimated utilizing the formula ‘volume = [(length + width)/2] x length x width’. Quantitation of the tumour volumes was analysed by Two-way ANOVA using GraphPad Prism9.

**Supplementary Figure 14.**
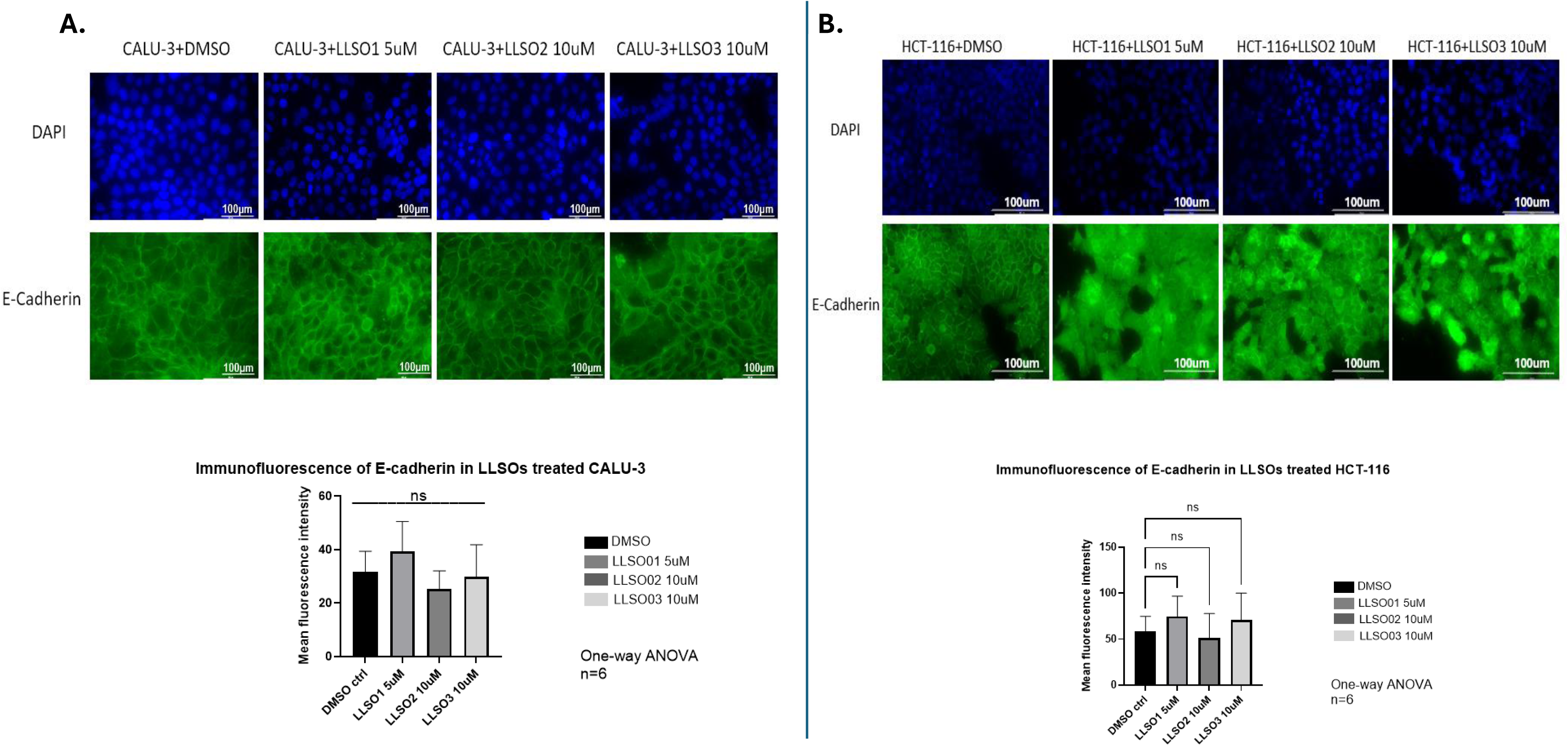
LLSOs have no effect on the expression of E-cadherin in CALU-3 and HCT-116 cells. Immunofluorescence analysis for E-cadherin expression was performed following 48 hours treatment of CALU-3 (**A**) or HCT-116 (**B**) cells with DMSO, 5μM LLSO1, 10μM LLSO2, and 10μM LLSO3. CALU-3 cells stained with mouse IgG were used as negative control and LNCaP cells were used as a positive control. None o f three LLSOs shows any impact on the E-cadherin expressing level. There are three wells repeats within each two independent experiments, n=6. Quantification of the E -cadherin signal is done with the ImageJ through using 9 fields of 3 repeat slides for each treatment. Photomicrographs were taken at 200x magnification; Scale bar is 100µm.

**Supplementary Figure 15.**
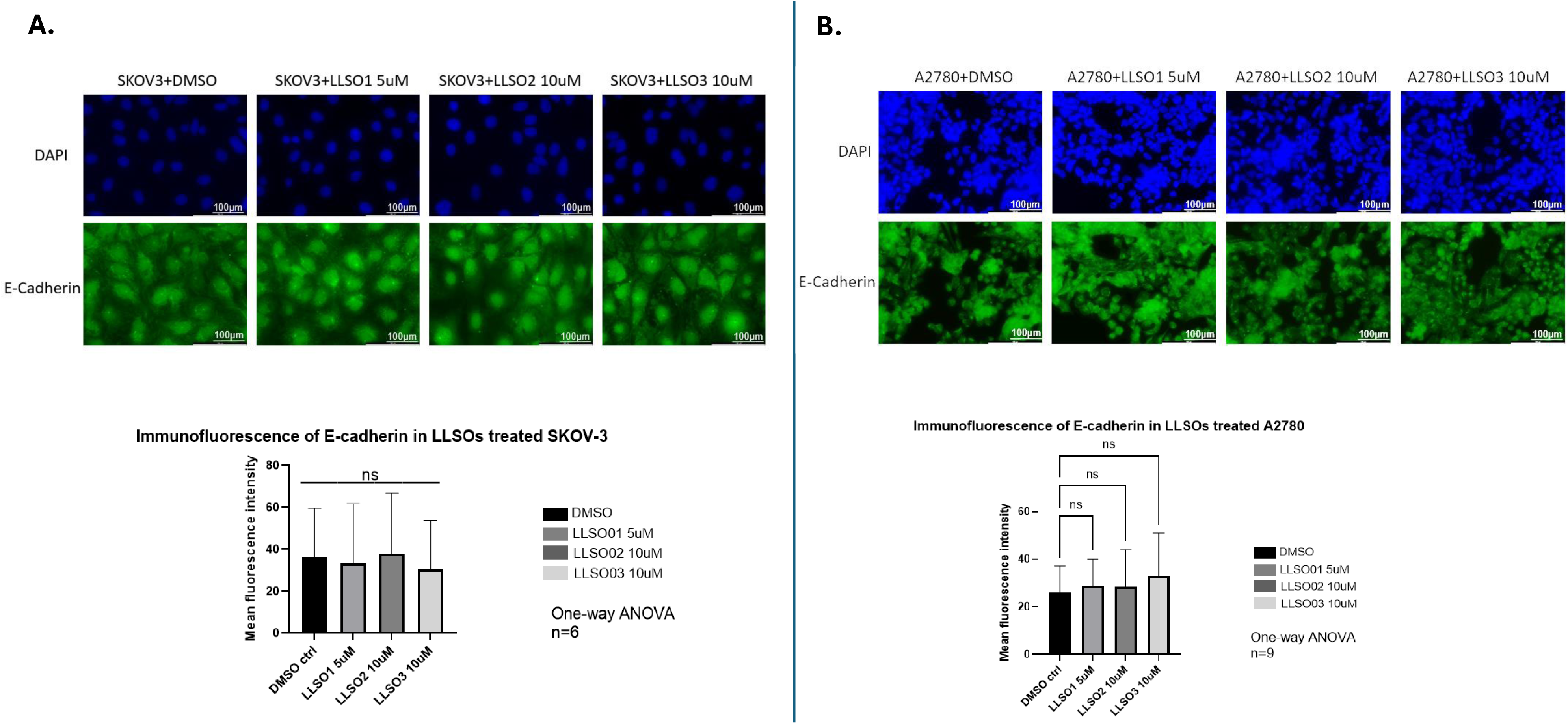
LLSOs have no effect on the expression of E-cadherin in SKOV-3 and A2780 cells. Immunofluorescence analysis for E-cadherin expression was performed following 48 hours treatment of SKOV-3 (**A**) or A2780 (**B**) cells with DMSO, 5μM LLSO1, 10μM LLSO2, and 10μM LLSO3. CALU-3 cells stained with mouse IgG were used as negative control and LNCaP cells were used as a positive control. None o f three LLSOs shows any impact on the E-cadherin expressing level. There are three wells repeats within each two independent experiments, n=6. Quantification of the E -cadherin signal is done with the ImageJ through using 9 fields of 3 repeat slides for each treatment. Photomicrographs were taken at 200x magnification; Scale bar is 100µm.

**Supplementary Figure 16.**
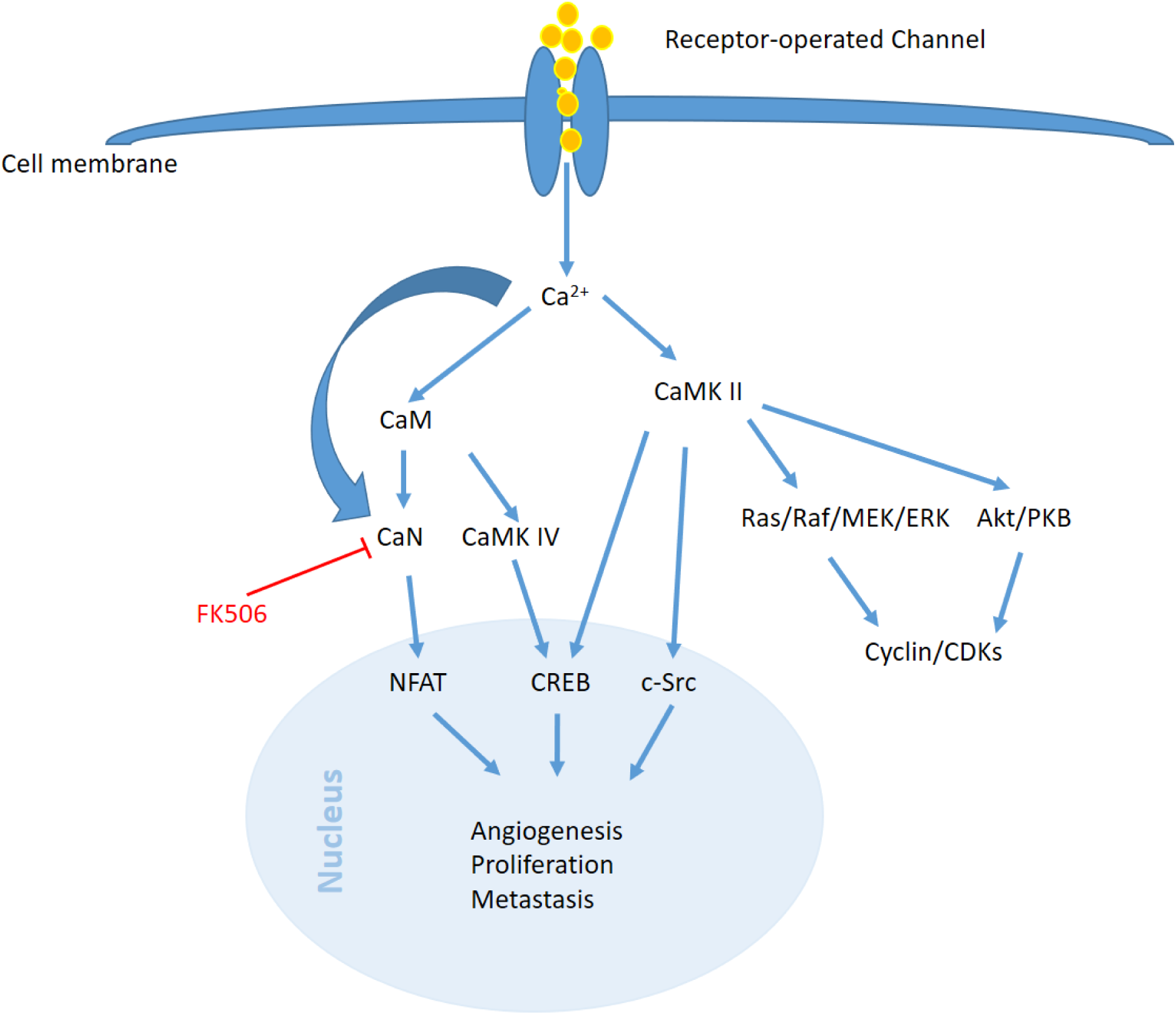
Hypothetical signaling pathway for LLSO1. LLSO1 inhibition of TTCC leads to the suppression of CaM, which subsequently inhibit the downstream CaMKII. Then this inhibition may potentially impede cancer progression by restricting the Ras/Raf/MEK/ERK pathway and the Akt/PKB pathway, both of which regulate the cyclin/cyclin-dependent kinase (CDK) complexes. Additionally, it is possible that c-Src can be inhibited by LLSO1 via the inhibition of CaMKII. LLSO1 might also inhibit CREB through inhibition of CaM via CaMKII and CaMKIV. Furthermore, NFAT, which is activated by CaN, can also be targeted by LLSO1 to impede EMT, since NFAT has been implicated in various aspects of cancer progression, such as apoptosis, proliferation, and invasion.

**Supplementary Figure 17.**
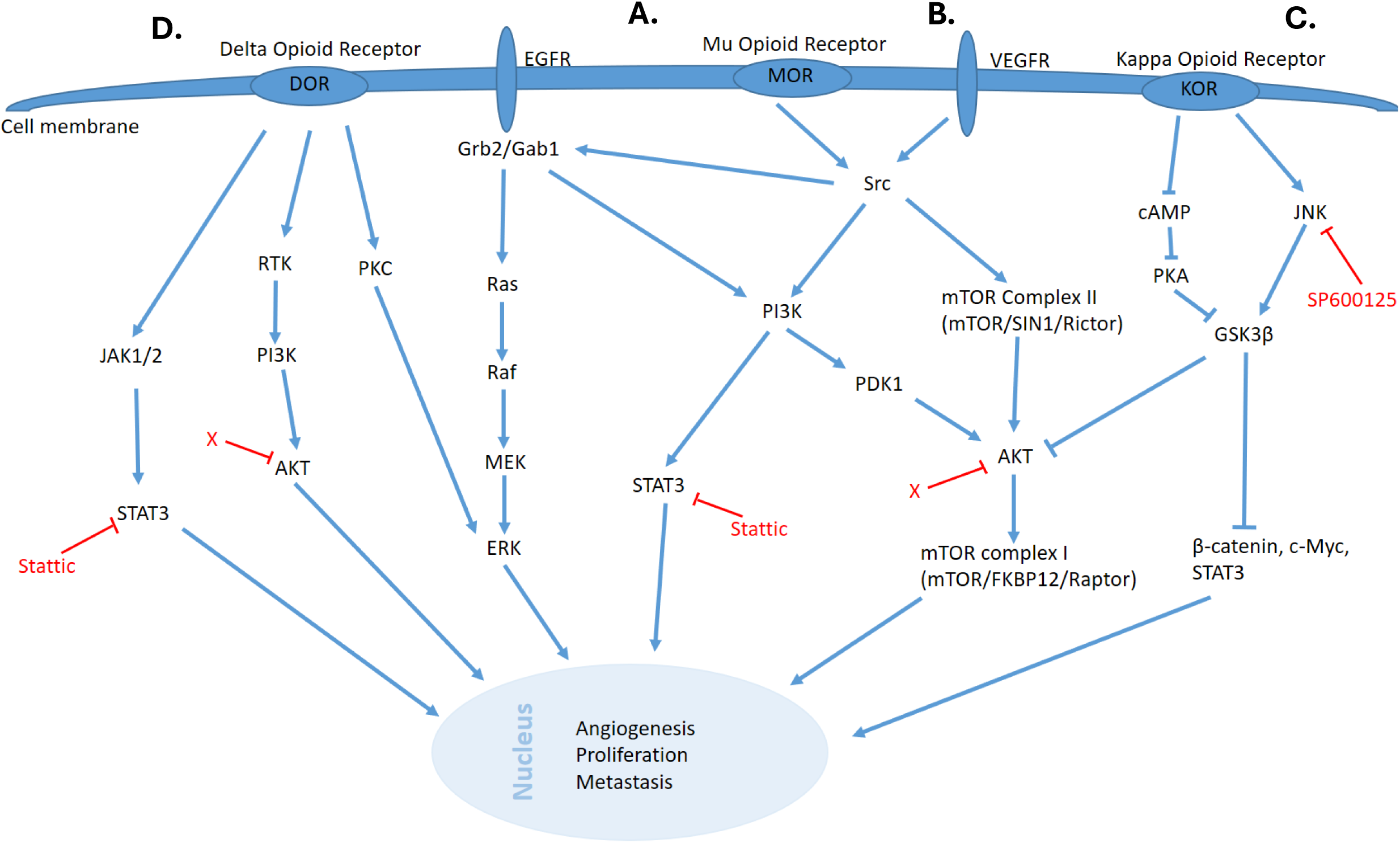
Hypothetical signaling pathway for LLSO3. **A.** LLSO3 may inhibit the MOR through blocking the transactivation of EGFR via Grb2/Gab1 and Scr suppression, as well as consequently inhibit the PI3K with downstream STAT3 and AKT activation. Meanwhile, either the interactions of CaM with MOR or the Ras-Raf complex activate MEK/ERK phosphorylation result in the EMT, proliferation and invasion of cells can also be targeted by LLSO3 (Lennon et al., 2014, Fujioka et al., 2011, Belcheva et al., 2001). **B.** LLSO3 might also inhibit MOR via VEGF and Src phosphorylation, and consequently through mTOR II formation and PI3K/PDK1 phosphorylation to block AKT followed by the mTOR I leading to the angiogenesis (Singleton et al., 2015). **C.** The effect of LLSO3 inhibition on KOR leads to activate of cAMP/PKA-mediated GSK3β inhibition, while simultaneously inhibiting JNK-mediated GSK3β activation. Thereby leading to the inhibition of c-Myc and β-catenin, which are typically overexpressed in cancer (Tian et al., 2006, Kuzumaki et al., 2012, Gao et al., 2014, Oren, 2003). **D.** LLSO3 might inhibit DOR through inhibition of the PKC/ERK, RTK/PI3K/AKT or JAK1/2-STAT3 pathway (Wei et al., 2016, Heiss, Ammer and Eisinger, 2009, Tripolt et al., 2021).

